# Lamb1a regulates atrial growth by limiting excessive, contractility-dependent second heart field addition during zebrafish heart development

**DOI:** 10.1101/2021.03.10.434727

**Authors:** Christopher J. Derrick, Eric J. G. Pollitt, Ashley Sanchez Sevilla Uruchurtu, Farah Hussein, Emily S. Noёl

## Abstract

During early vertebrate heart development, the heart transitions from a linear tube to a complex asymmetric structure. This process includes looping of the tube and ballooning of the emerging cardiac chambers, which occur simultaneously with growth of the heart. A key driver of cardiac growth is deployment of cells from the Second Heart Field (SHF) into both poles of the heart, with cardiac morphogenesis and growth intimately linked in heart development. Laminin is a core component of extracellular matrix (ECM) basement membranes, and although mutations in specific laminin subunits are linked with a variety of cardiac abnormalities, including congenital heart disease and dilated cardiomyopathy, no role for laminin has been identified in early vertebrate heart morphogenesis. We identified dynamic, tissue-specific expression of laminin subunit genes in the developing zebrafish heart, supporting a role for laminins in heart morphogenesis. *lamb1a* mutants exhibit cardiomegaly from 2dpf onwards, with subsequent progressive defects in cardiac morphogenesis characterised by a failure of the chambers to compact around the developing atrioventricular canal. We show that loss of *lamb1a* results in excess addition of SHF cells to the atrium, revealing that Lamb1a functions to limit heart size during cardiac development by restricting SHF addition to the venous pole. *lamb1a* mutants exhibit hallmarks of altered haemodynamics, and specifically blocking cardiac contractility in *lamb1a* mutants rescues heart size and atrial SHF addition. Furthermore, we identify that FGF and RA signalling, two conserved pathways promoting SHF addition, are regulated by heart contractility and are dysregulated in *lamb1a* mutants, suggesting that laminin mediates interactions between SHF deployment, heart biomechanics, and biochemical signalling during heart development. Together, this describes the first requirement for laminins in early vertebrate heart morphogenesis, reinforcing the importance of specialised ECM composition in cardiac development.

## Introduction

Tissue morphogenesis requires tight coordination of changes in cell shape and organisation, gene expression, and tissue patterning, together with the integration of both intrinsic and extrinsic signalling cues. Cardiac development represents an excellent example of such complex tissue morphogenesis, where the linear heart tube undergoes growth, local tissue deformation, and functional regionalisation, all of which must be coordinated between multiple tissue layers. This early heart morphogenesis is required to ensure subsequent events in cardiac development, such as proper alignment of the chambers to facilitate septation, chamber ballooning, and connection to the rest of the cardiovascular system, proceed normally. The importance and complexity of robust heart morphogenesis are evident in the prevalence of congenital heart diseases (CHDs), which occur in at least 1% of live births and are the leading cause of birth defect related deaths worldwide [1,2].

Heart looping and chamber ballooning are key stages in cardiac development during which the heart tube undergoes a complex morphological rearrangement resulting in a helical looped tube in mouse, and a planar looped tube in zebrafish [3]. Concomitant with looping and ballooning morphogenesis, multiple processes further shape the organ, including a dramatic increase in myocardial cell number, primarily achieved through addition of cells to the developing heart from a conserved pool of progenitor cells termed the second heart field (SHF) [4]. The SHF is located in the mesoderm adjacent to the developing heart, and is contiguous with the poles of the myocardial heart tube. During cardiogenesis in mouse and chick, SHF addition generates a significant proportion of cardiac tissue, giving rise to the right ventricle, atria, and inflow and outflow tracts [5–8]. Similarly, in zebrafish the SHF makes a major contribution to the inflow tract at the base of the atrium, the single ventricle, and the outflow tract (OFT) [9–13]. The signaling pathways required for SHF addition are highly conserved [14,15]. *Fgf8* (*fgf8a* in zebrafish) signalling promotes SHF addition, whilst an opposing Retinoic Acid (RA) gradient limits SHF addition to the arterial pole [16–18]. Downstream, SHF addition to both poles of the heart is dependent on the transcription factor *Isl1* in mice [19], whilst *isl1a* is required only for venous pole addition in zebrafish with *isl2b* driving arterial SHF addition [13,20]. SHF addition and cardiac morphogenesis are tightly linked, with defects in SHF addition leading to heart malformations and congenital heart disease [21].

Mechanical forces have also been linked with SHF addition. Recent studies have demonstrated that the SHF epithelium is under tension, which is proposed to regulate cell orientation and cell division, driving the extension of the linear heart tube [22], and it has been suggested that heart tube contractility could contribute to this tension. Furthermore, Wnt5a-mediated changes in cell-cell adhesion are responsible for regionalised differences in pushing/pulling forces in the outflow tract that are necessary for SHF deployment [23]. Heart function begins prior to completion of heart morphogenesis and thus cardiomyocyte contractility, blood flow, SHF addition, and morphogenesis are also coupled during heart development. This is highlighted by a failure in atrioventricular valve development [24–28] and failure in initiation of trabeculation [29–31] in zebrafish lacking the cardiac Troponin *tnnt2a*, which is required for heart contractility [28]. Independent from cardiomyocyte contractility, the sensation of blood flow is also required for heart morphogenesis. The forces generated by blood flow results in localised induction of transcription factors such as *klf2a* which functions in the endocardium to promote regionalised cell shape changes [32] and proliferation associated with early valve development [24,25]. Thus a complex interplay of biochemical and biomechanical cues are required for robust heart morphogenesis during development.

In the developing heart the myocardium and endocardium are separated by a layer of extracellular matrix (ECM) termed the cardiac jelly. The ECM is an important signalling centre which influences biochemical signalling between cells, as well as providing biomechanical stimuli. Numerous studies have highlighted the importance of the cardiac jelly during development of the heart [33] such as regionalised ECM synthesis driven by *klf2a* [34,35], or balanced ECM synthesis and degradation during trabeculation [36,37]; however, comparatively little is known about the specific roles that individual ECM components play in promoting heart looping, chamber morphogenesis, and cardiac growth.

Laminins are large heterotrimeric complexes laid down early during ECM deposition. Consisting of an alpha, beta and gamma chain, laminin trimers are an essential component of the basement membrane where they polymerise into a lattice network, interacting with integrin receptors on the cell membrane and facilitating the organisation of higher order ECM structures in the interstitial matrix [38,39]. Multiple alpha, beta, and gamma subunits are encoded in the genome, and can assemble into a variety of trimer isoforms which often exhibit tissue restricted expression, and play specific roles in different tissue contexts [40]. Unsurprisingly given the central role for laminins in coordinating ECM composition and cell-ECM signalling, there is clear evidence that laminins play important roles in human heart development and function. Deleterious mutations in *LAMA4* have been identified in patients with dilated cardiomyopathy, accompanied by defects in endothelial cell attachment [41]. Further analysis of heart development upon combinatorial knockdown of zebrafish *lama4* and *integrin-linked kinase (ilk)* suggest the laminin-integrin signalling axis promotes endocardial integrity [41]. A mutation in *LAMA5* has been associated with a multi-systemic disorder which includes cardiac abnormalities [42], and ~30% of patients with Dandy-Walker Syndrome, a rare brain malformation linked to mutations in *LAMC1*, also present with CHDs [43,44]. Whilst these studies highlight an important requirement for laminins in cardiac form and function, and despite the fundamental role laminins play in ECM organisation and matrix-cell signalling, little is known about how they support heart development.

Current vertebrate models have provided limited insights into the role of laminins in cardiac development and function. *Lama4* mutant mice survive postpartum without overt cardiac defects, although the pups are haemorrhagic at birth and over time exhibit hallmarks of sustained hypoxia, developing enlarged hearts and larger cardiomyocytes [45,46]. Functional analyses reveal that whilst loss of *Lama4* does not affect heart rate, *Lama4* mutants display cardiac dysfunction including arrhythmia and reduced left ventricular end diastolic diameter [46], alongside increased frequency of sudden death [45], suggesting a conserved role in endothelial integrity and additional functions in cardiomyocyte biology. While the *Lama4* mutant mouse facilitates mechanistic understanding of how laminins may regulate endothelial function, interrogating the role of laminins more broadly in heart development is less straightforward due to early lethality in both *Lamc1* and *Lamb1* mutant mice [47,48]. Direct evidence supporting a role for laminins in heart development comes from *Drosophila*, where laminins promote the formation and integrity of the dorsal vessel [49,50]. A further study in zebrafish demonstrated that the extraembryonic yolk syncytial layer facilitates cardiomyocyte migration by regulating *fibronectin 1a* expression and laminin deposition, supporting a conserved requirement for laminins in vertebrate heart development [51]. However, vertebrate studies are yet to define the key requirements for laminins in early cardiac morphogenesis.

Here we identify tissue-specific expression of multiple laminin subunits in the developing zebrafish heart during early cardiac morphogenesis. Through targeted mutagenesis of the highly conserved laminin genes *lamc1* and *lamb1a*, we identify two novel requirements for laminins during heart development: promoting heart looping morphogenesis and restricting cardiac size. We show that Lamb1a controls atrial growth by limiting SHF addition to the venous pole, and demonstrate that the excessive atrial SHF addition in *lamb1a* mutants can be rescued by blocking heart contractility. Finally, we demonstrate that loss of either contractility or *lamb1a* disrupts expression of FGF and RA-responsive genes in the heart, supporting a role for laminins in coupling mechanical force, intercellular signalling, and cardiac growth. Together, this study presents the first reported role for laminins in the early development of the vertebrate heart and highlights a crucial requirement for the ECM in coordinating distinct aspects of tissue morphogenesis.

## Results

### Laminins display dynamic, tissue-specific expression during early zebrafish heart morphogenesis

To investigate the role of laminin complexes in early vertebrate heart morphogenesis we probed a previously-published transcriptomic analysis of cardiac gene expression to identify laminin subunit genes expressed in the heart tube [52], and in combination with an *in situ* hybridization screen identified a subset of laminin subunits with cardiac expression during early stages of heart looping (Fig 1, Supplemental Fig S1).

**Figure 1.**
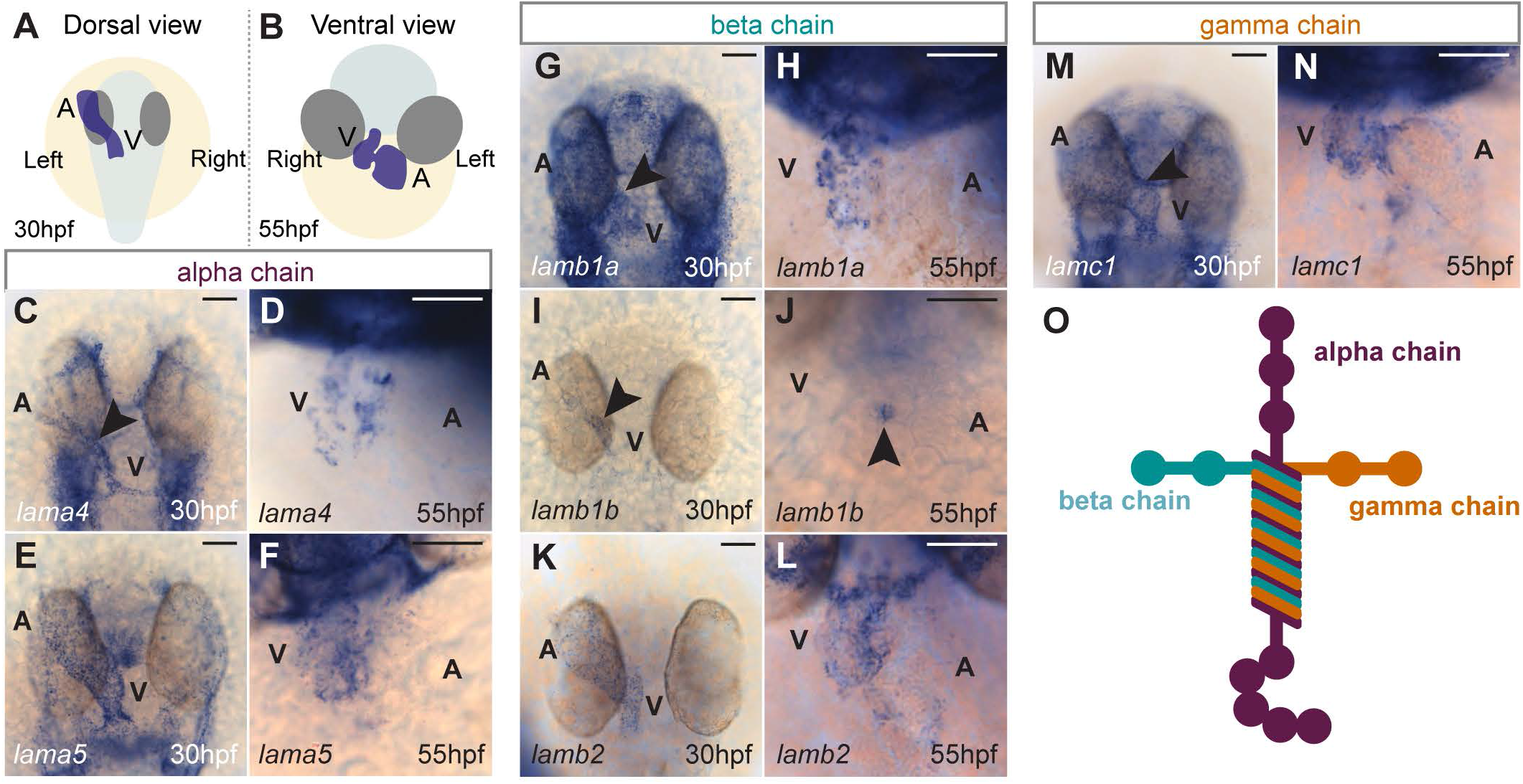
Dynamic expression of laminin subunit genes during heart morphogenesis. A,B: Schematic depicting position of heart (blue) in a 30hpf zebrafish embryo (dorsal view) and a 55hpf zebrafish embryo (ventral view). C-F: mRNA *in situ* hybridization expression analysis of laminin alpha chain subunits *lama4* (C,D) and *lama5* (E,F) in the heart. G-L: mRNA *in situ* hybridization expression analysis of laminin beta subunit chains *lamb1a* (G,H), *lamb1b* (I,J) and *lamb2* (K,L) in the heart. M-N: mRNA *in situ* hybridization expression analysis of gamma subunit *lamc1* in the heart. Anterior to top. V - Ventricle, A - Atrium. Scale bars: 50μm. O: Schematic depicting laminin subunit organisation into heterotrimeric structure.

Zebrafish share ten of the twelve mammalian laminin subunits, with no orthologs for *lamb3* or *lamc2* and possess two duplicated beta subunits - *lamb2l* and *lamb1b* [53]. At 30hpf (hours post fertilisation, Fig 1A), during early heart tube morphogenesis, six laminin subunits are expressed in the zebrafish heart: two alpha chains, *lama4* and *lama5*, (Fig 1C,E); three beta chains, *lamb1a, lamb1b*, and *lamb2* (Fig 1G,I,K) and a single gamma chain: *lamc1* (Fig 1M). Since specific laminin isoforms can exhibit tissue-specific deposition, we carried out two colour fluorescent *in situ* hybridization at 30hpf to identify whether the myocardium and endocardium have a specific laminin expression profile (Supplemental Fig S1). This identified two endocardial laminin subunit genes: *lama4* and *lamb1b* (Supplemental Fig S1A-B), two myocardial laminins: *lama5* and *lamb2* (Supplemental Fig S1C-D) and two laminin subunits that are expressed in both the myocardium and endocardium: *lamb1a* and *lamc1* (Supplemental Fig S1E-F).

While at 30hpf the majority of laminin genes are expressed along the length of the heart tube, following initial heart looping morphogenesis at 55hpf (Fig 1B), the expression of most laminin subunits has become restricted to the ventricle and atrioventricular canal (Fig 1D,F,H,L,N) with the exception of *lamb1b* which is expressed only in the atrioventricular canal (Fig 1J). This dynamic spatiotemporal control of specific laminin subunit expression during early heart development suggests that individual endocardial or myocardial-derived laminin complexes may drive early heart tube morphogenesis.

### lamc1 and lamb1a regulate heart morphology and size during cardiac development

Having identified multiple potential laminin complexes expressed in the heart during early morphogenesis, we wanted to examine the role of laminins during heart development. Laminins are a heterotrimeric complex of a single alpha, beta and gamma chain (Fig 1O) which are assembled in the cell prior to deposition into the ECM, thus the removal of a single subunit is sufficient to prevent secretion of the complex from the cell [54]. Therefore, to investigate the requirement for laminins during heart looping morphogenesis we targeted the single gamma subunit *lamc1* (the homolog of human *LAMC1*), expressed in both the myocardium and endocardium (Fig 1M-N and Supplemental Fig S1F).

To first investigate the role of *lamc1* in zebrafish heart development we performed CRISPR-Cas9 mutagenesis, injecting two gRNAs targeting *lamc1* together with Cas9 protein into 1-cell stage wild-type embryos to generate transient F0 *lamc1* mutants (crispants) [55]. *lamc1* crispants recapitulate the morphological phenotype of stable *lamc1* mutants with a high level of efficacy, whilst uninjected embryos or injection controls (gRNA only or Cas9 only) are morphologically normal (Supplemental Fig S2A-D) [56–58]. Despite a profoundly shortened body axis, *lamc1* crispants formed a beating heart tube by 26hpf indicating laminin is not required for initial heart tube formation. At 2dpf *lamc1* mutants exhibit mild pericardial oedema, suggesting possible defects in heart looping morphogenesis (Supplemental figure S2D). We therefore assessed the impact of loss of Lamc1 function on heart morphology by mRNA *in situ* hybridization analysis of *myl7* expression (Fig 2A-D), quantifying looping ratio and heart size at 55hpf and 72hpf after initial heart looping morphogenesis has occurred (Fig 2E-H, Supplemental Fig S2E-F) [52]. At 55hpf, *lamc1* F0 crispants have failed to undergo correct heart looping morphogenesis (Fig 2A-B), displaying a significant reduction in heart looping ratio compared to control embryos (Fig 2E), although the size of the heart appears relatively normal (Fig 2F). By 72hpf, in addition to abnormal cardiac morphology (Fig 2D, and Supplemental Fig S2E) *lamc1* F0 crispant hearts also appear significantly larger than in control embryos (Fig 2H, Supplemental Fig 2F). This demonstrates that laminins promote initial heart looping morphogenesis and may subsequently function to restrict cardiac size.

**Figure 2.**
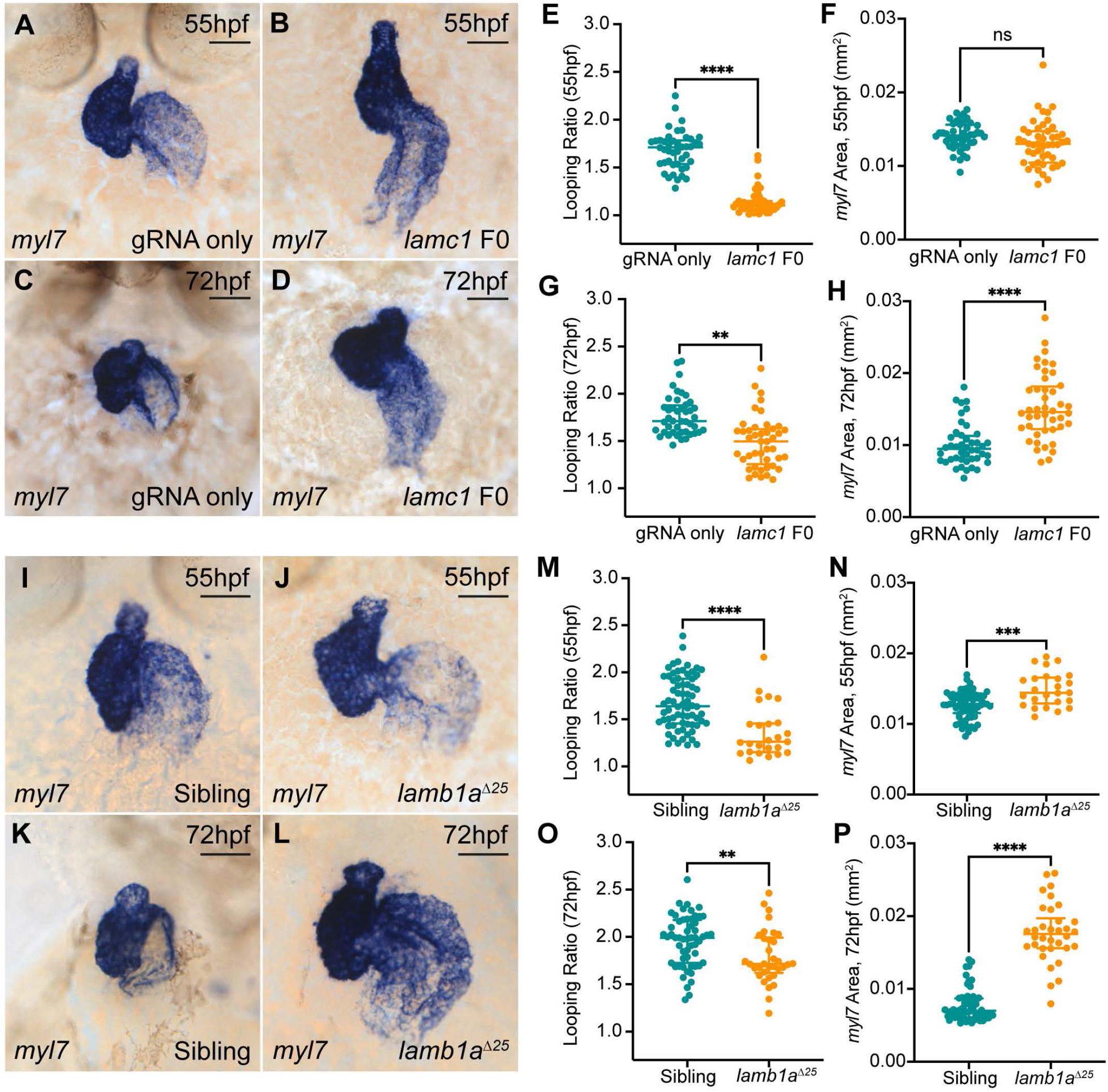
Laminins perform multiple roles during zebrafish heart morphogenesis. A-D: mRNA *in situ* hybridization analysis of *myl7* expression in control embryos injected with *lamc1*-targeting gRNAs only (A,C) or with *lamc1*-targeting gRNAs together with Cas9 protein (*Lamc1 F0*, B-D) at 55hpf and 72hpf. E-H: Quantitative analysis of looping ratio (E, G) and *myl7* area (F,H) in gRNA-injected controls (55hpf: n=44; 72hpf: n=44) and *lamc1* F0 crispants (55hpf: n=47; 72hpf: n=44). *lamc1* crispants exhibit reduced heart looping at 55hpf and 72hpf, and an increased area of *myl7* expression at 72hpf. Horizontal bars indicate median with interquartile range, comparative statistics performed using Kruskal-Wallis test. I-L: mRNA *in situ* hybridization analysis of *myl7* expression in sibling (I,K) and *lamb1a^Δ25^* mutants (J,L) at 55hpf and 72hpf. M-P: Quantitative analysis of looping ratio (M,O) and *myl7* area (N,P) in sibling (55hpf: n=70; 72hpf: n=56) and *lamb1a^Δ25^* mutant embryos (55hpf: n=25; 72hpf: n=34). *lamb1a^Δ25^* mutants exhibit a mild reduction in heart looping at 55hpf, and an increased area of *myl7* expression at 55hpf and 72hpf. Scale bars: 50μm. Comparative statistics performed using Mann Whitney test, **** = *p* < 0.0001, *** = p < 0.001, ** = p < 0.01, * = p< 0.05, ns = not significant.

Distinct laminin complexes play varied yet specific roles in different developmental contexts [40]. Since *lamb1a* (the homolog of human *LAMB1*) exhibits similar expression dynamics and tissue-specificity as *lamc1* (Fig 1G,H,M,N), this suggested that Lamb1a and Lamc1 may together form part of the specific laminin complexes required for heart development. To confirm this role for *lamb1a* we generated two stable mutant alleles *lamb1a^Δ19^* and *lamb1a^Δ25^* using CRISPR-Cas9 mediated genome editing (Supplemental Fig S2G-M). At 55hpf *lamb1a* mutants display a much less severe heart looping defect when compared to *lamc1* crispants (Fig 2B,J), but instead present a mild increase in heart size (Fig 2N, Fig S2N,O,R). Strikingly, loss of *lamb1a* results in progressive defects in cardiac size, characterised by significantly larger hearts compared to siblings at 72hpf (Fig 2K,L,P, Supplemental Fig S2P,Q,S). While mutant alleles for both *lamc1* and *lamb1a* have been previously described [56,57], defects in heart development have not been reported, although in one study heart development in *lamb1a* mutants was examined but only prior to 35hpf [59]. Analysis of heart morphology in *grumpy^tj299a^* (*gup, lamb1a*) and *sleepy^sa379^* (*sly, lamc1*) mutants revealed identical phenotypes to our novel loss of function models (Supplemental. Fig S3). Together, these data demonstrate two previously uncharacterised requirements for laminins in vertebrate heart morphogenesis, where Lamc1-containing trimers promote heart looping and restrict cardiac size, while Lamb1a-containing trimers primarily function to restrict heart size.

The difference in phenotypes between *lamb1a* and *lamc1* mutants suggested other laminin beta subunits may act during early heart morphogenesis, or functionally compensate for loss of *lamb1a.* We examined the impact of loss of *lamb1a* on the expression of the two other identified laminin beta subunits (*lamb1b* and *lamb2*) in the heart at 30hpf (Supplemental Fig S4A-D). At 30hpf a striking upregulation and expansion of *lamb1b* expression was observed in *lamb1a^Δ25^* mutants (Supplemental Fig S4A-B), whilst *lamb2* expression was unaffected (Supplemental Fig S4C-D). This suggested that *lamb1b* could compensate for loss of *lamb1a*, resulting in the weaker looping morphogenesis phenotype in *lamb1a* mutants when compared to loss of *lamc1*. To investigate this we used CRISPR-Cas9-mediated genome editing to generate two *lamb1b* promoter deletion alleles: *lamb1b^Δ183^* and *lamb1b^Δ428^* (Supplemental Fig S4E-F). We generated *lamb1b;lamb1a* double heterozygous adult fish, incrossed them to obtain *Lamb1b;lamb1a^Δ25^* double mutant embryos and examined *Lamb1b* expression at 30hpf by *in situ* hybridization to confirm the absence of *Lamb1b* transcript (Supplemental Fig S4G-J). Analysis of heart size and morphology in *Lamb1b;lamb1a^Δ25^* double mutants at 55hpf revealed that loss of *Lamb1b* did not modify the *lamb1a* mutant phenotype, demonstrating that despite its upregulation in *lamb1a^Δ25^* mutants, does not compensate for loss of *lamb1a* (Supplemental Fig S4K-N). This further supports the hypothesis that *lamc1* and *lamb1a* play distinct roles in cardiac development.

### lamb1a limits SHF addition to the venous pole

We have identified a requirement for laminins in driving looping morphogenesis and restricting organ size during heart development (Fig 2). These two processes are uncoupled in the *lamb1a* mutant, an interesting finding as often defects in heart size are coupled with a severe impact on looping morphology, such as the loss of Cerebral Cavernous Malformation (CCM) pathway components where cardiac chambers are larger but morphology is also severely disrupted [60–62]. The *lamb1a* mutant thus represents an excellent model to investigate the mechanism by which laminins limit heart growth independent of the morphogenesis of the tissue.

To determine whether growth of a specific chamber was particularly impacted by loss of *lamb1a* we examined chamber size at 55hpf and 72hpf by *in situ* hybridization analysis of *myl7l* and *myh6* expression in the ventricle and atrium respectively. *lamb1a* mutants display a significant increase in the size of both chambers (Fig 3A-F) with progressive enlargement between 55hpf and 72hpf, suggesting that laminin normally limits growth of both chambers. Increase in cardiac size could be due to either cardiomyocyte hypertrophy, or increased cell number. Loss of *lama4* and *integrin-linked kinase* (the intracellular effector of laminin-integrin signalling) have previously been associated with dilated cardiomyopathy [41], with zebrafish *ilk* mutants exhibiting stretched cardiomyocyte morphology. This suggested that cardiomegaly in *lamb1a* mutants may be due to enlarged cardiomyocytes. We quantified internuclear distance in both chambers in wild type and *lamb1a^Δ25^* mutant *Tg*(*-5.1myl7:DsRed2-NLS*) embryos (henceforth *Tg*(*myl7:DsRed*)), in which myocardial nuclei express DsRed2, at 55hpf and 72hpf (Fig 3G-I). In contrast to our expectation that loss of *lamb1a* would result in enlarged cardiomyocytes, *lamb1a^Δ25^* mutant embryos do not exhibit increased internuclear distance when compared to siblings, demonstrating that Lamb1a is not restricting cardiomyocyte size (Fig 3J,K). Since cardiomyocytes are not enlarged in *lamb1a^Δ25^* mutants, we therefore hypothesised that cardiomegaly in *lamb1a* mutants results from increased cell number, and quantified DsRed-positive cardiomyocytes in sibling and *lamb1a^Δ25^* mutant embryos at 55hpf and 72hpf. We observed no profound increase in cell number in the ventricle at either stage; however, we did find a significant increase in the number of atrial cells in *lamb1a^Δ25^* mutant embryos at both 55hpf and 72hpf when compared to sibling embryos (Fig 3L,M). Together this suggests that Lamb1a controls atrial size through regulating cell number.

**Figure 3.**
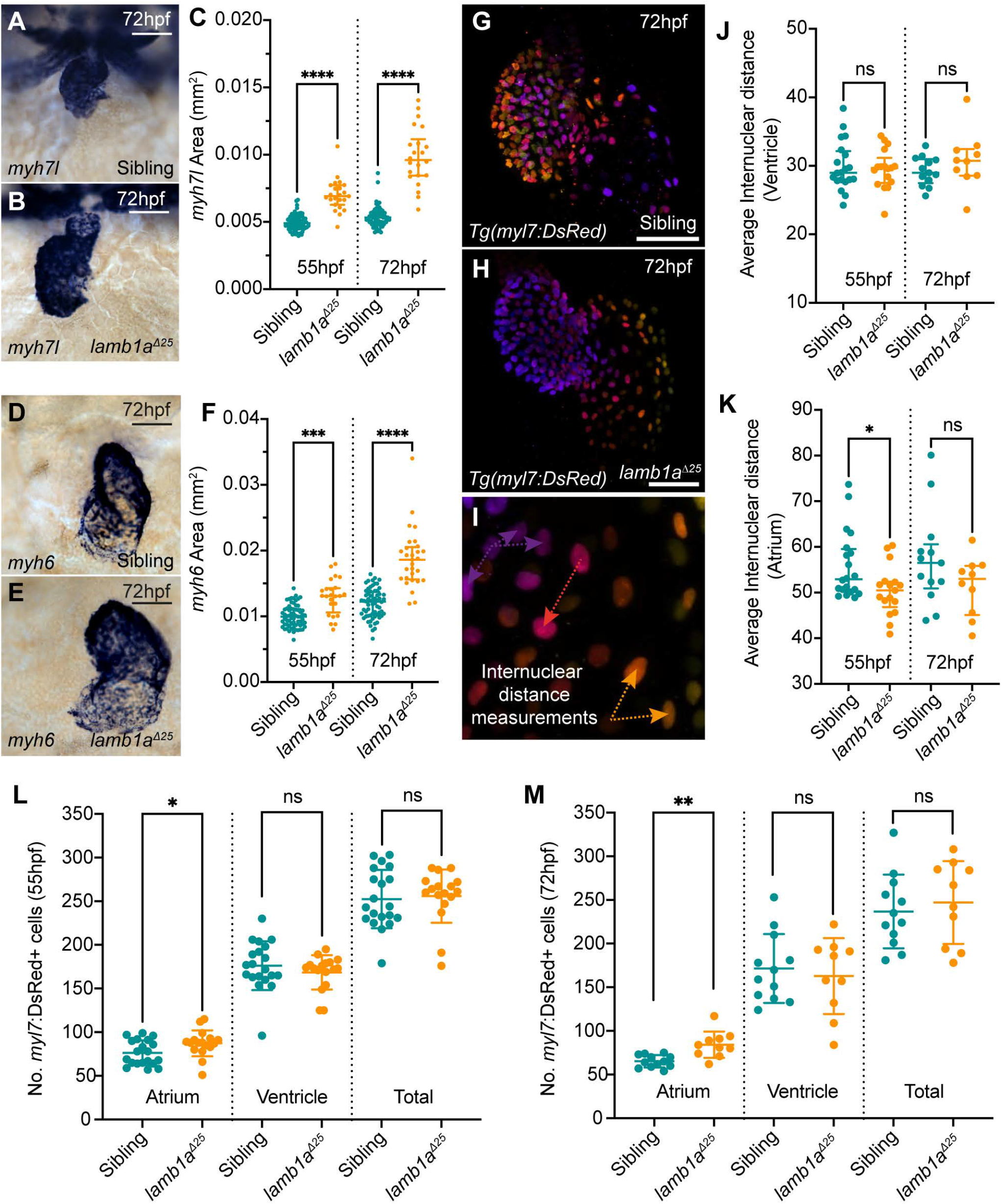
lamb1a mutants have increased atrial cells. A-B: mRNA *in situ* hybridization analysis of *myh7l* expression in the ventricle of sibling (A) and *lamb1a^Δ25^* mutant embryos (B) at 72hpf. C: Quantification of *myh7l* expression area in sibling (55hpf: n=72; 72hpf: n=67) and *lamb1a^Δ25^* mutants (55hpf: n=23; 72hpf: n=22). D-E: mRNA *in situ* hybridization analysis of *myh6* expression in the atrium of sibling (D) and *lamb1a^Δ25^* mutant embryos (E) at 72hpf. F: Quantification of *myh6* expression area in sibling (55hpf: n=65; 72hpf: n=64) and *lamb1a^Δ25^* mutants (55hpf: n=24; 72hpf: n=29). Horizontal bars indicate median with interquartile range, comparative statistics performed using Kruskal-Wallis test. G-I: Depth-coded maximum intensity projections of confocal image z-stacks in *Tg(myl7:DsRed)* transgenic sibling (G) and *lamb1a^Δ25^* mutant embryos (H) at 72hpf. Internuclear distance is quantified between nuclei on the same face of the heart which occupy similar z-positions (I). J-K: Quantification of average internuclear distance at 55hpf and 72hpf in the ventricle (J) and atrium (K) of sibling (55hpf: n=20; 72hpf: n=17) and *lamb1a^Δ25^* mutant embryos (55hpf: n=13; 72hpf: n=10), demonstrating a mild decrease in the internuclear distance of *lamb1a^Δ25^* mutant atrial cells at 55hpf (K). Horizontal bars indicate median with interquartile range, comparative statistics performed using Brown-Forsythe and Welch ANOVA with multiple comparisons. L-M: Quantification of DsRed+ cells in the myocardium of *Tg(myl7:DsRed)* transgenic sibling (55hpf: n=20; 72hpf: n=17) and *lamb1a^Δ25^* mutant embryos (55hpf: n=13; 72hpf: n=10) at 55hpf (L) and 72hpf (M). *lamb1a^Δ25^* mutants have a significant increase in atrial cell number at both stages. Scale bars: 50μm. Chamber-specific analyses - unpaired t-test with Welch’s correction. All statistical analyses: **** = p < 0.0001, *** = p < 0.001, ** = p < 0.01, * = p< 0.05, ns = not significant.

During morphogenesis, the heart grows primarily through addition of cells to the poles of the heart from the second heart field (SHF), a conserved population of cells situated in the mesoderm adjacent to the heart. While previous studies have demonstrated that SHF addition to the arterial pole of the heart is sensitive to perturbations in ECM composition [33], comparatively less is known about SHF addition to the venous pole, and in particular how the ECM may regulate this process. Since *lamb1a^Δ25^* mutants exhibit increased atrial cell number in the heart during the window of SHF addition, we hypothesised that loss of Lamb1a-containing laminin trimers leads to increased cardiac size through elevated SHF addition. We visualised SHF addition in *Tg(myl7:eGFP);Tg(myl7:DsRed)* double transgenic sibling and *lamb1a^Δ25^* mutant embryos, where cardiomyocytes derived from the original linear heart tube/first heart field are marked by both GFP and DsRed expression, whilst cells recently added from the SHF are GFP-positive only [13] (Fig 4C,C’). At 55hpf, *lamb1a^Δ25^* mutants appeared to have a larger GFP+/DsRed-area at the venous pole of the heart, suggesting an increased number of SHF cells (Fig 4A-B”). Quantification of GFP+/DsRed-cell number in the atrium revealed a significant increase in SHF cells at the venous pole of *lamb1a^Δ25^* mutant embryos compared to siblings (Fig 4D,E), demonstrating that Lamb1a limits atrial size by restricting SHF addition to the venous pole.

**Figure 4.**
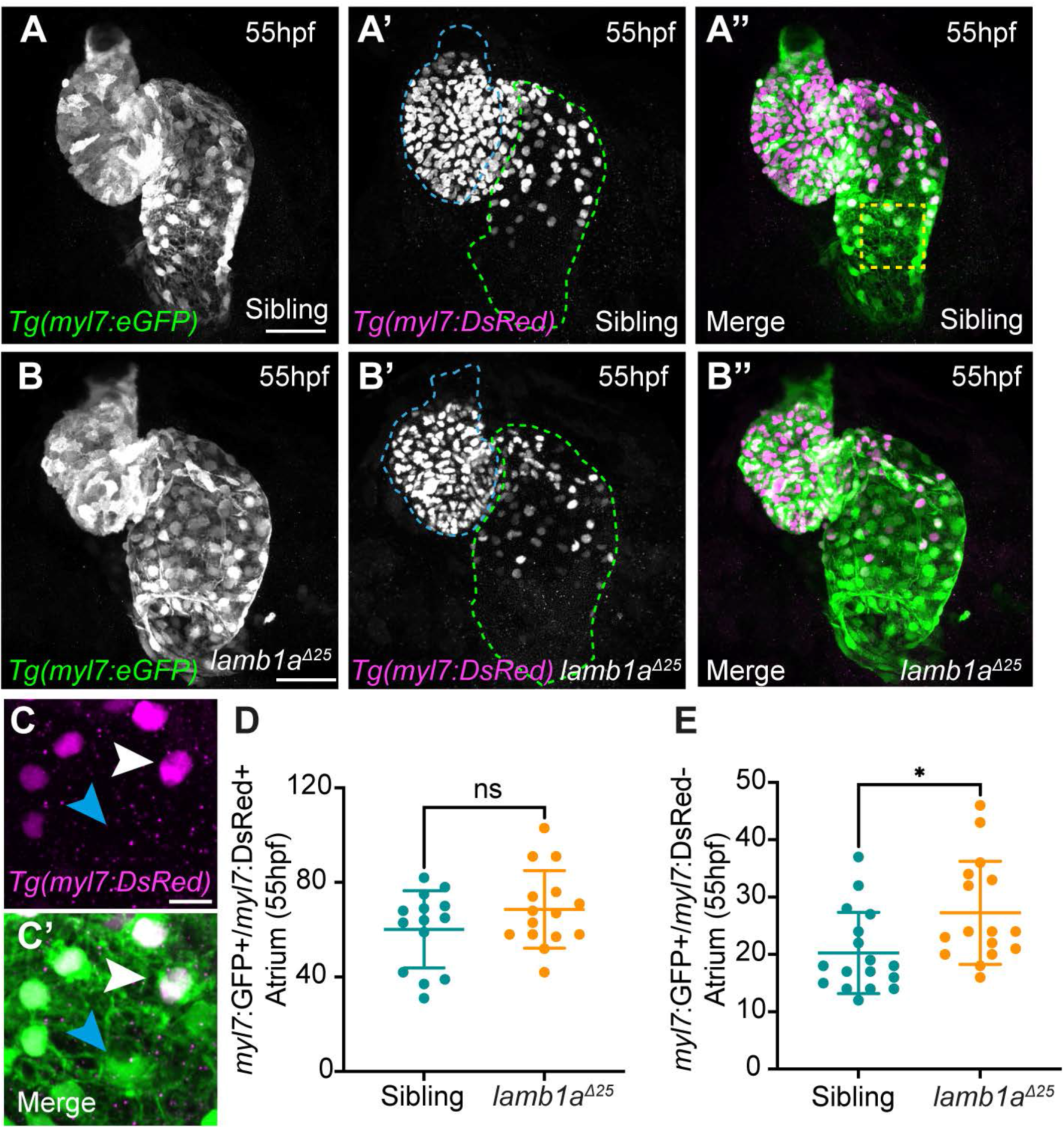
lamb1a limits SHF addition to the venous pole. A-B’’: Maximum intensity projections of confocal image z-stacks in *Tg(myl7:eGFP);Tg(myl7:DsRed)* double transgenic sibling (A-A’’) and *lamb1a^Δ25^* mutant embryos (B-B’’) at 55hpf. GFP+/DsRed-SHF cells can be visualised at both the venous (green dotted line) and arterial (blue dotted line) poles of the heart. C-C’: Higher magnification of the boxed area in A’’. Double GFP+/DsRed+ cells represent ‘older’ cardiomyocytes (white arrowhead), while GFP+/DsRed-cells represent newly added SHF cells (blue arrowhead). D-E: Quantification of double GFP+/DsRed+ cardiomyocytes (D) and GFP+/DsRed-SHF cells (E) in sibling (n=17) and *lamb1a^Δ25^* mutant embryos (n=16) at 55hpf reveals an increase in newly-added SHF cells in *lamb1a^Δ25^* mutants compared to siblings. Scale bars A-B: 50μm, C: 10μm. Horizontal bars indicate mean +/- SD, comparative statistics performed using Kolmogorov-Smirnov test, * = p < 0.05, ns = not significant.

The elevated SHF addition to the venous pole of *lamb1a^Δ25^* mutants could be due to an increased SHF progenitor pool. We examined expression of the ECM component *ltbp3*, and the transcription factor *isl1a*, which are expressed in SHF cells and are required in zebrafish for SHF addition to the arterial and venous poles respectively [10,13]. At 30hpf, no differences in expression domain or levels of either *ltbp3* or *isl1a* were observed in *lamb1a^Δ25^* mutant embryos when compared to siblings (Supplemental Fig S5A-D). At 55hpf while the level of *ltbp3* expression appears mildly reduced in the outflow tract of *lamb1a^Δ25^* mutants, the domain of expression does not appear to be expanded (Supplemental Fig S5E-F). Furthermore, *isl1a* expression at the venous pole of *lamb1a^Δ25^* mutant hearts is comparable with siblings at 55hpf (Supplemental Fig S5G-H’). Together this suggests that the increase in SHF addition in *lamb1a^Δ25^* mutants is not due to enlargement of the SHF domain.

### Excessive second heart field addition to the venous pole in lamb1a mutants is dependent on heart contractility

Our finding that loss of *lamb1a* leads to increased SHF addition to the atrium without expansion of the SHF domain suggested a SHF specification-independent mechanism of altered SHF addition. Our previous findings that *lamb1b* upregulation in *lamb1a* mutants does not appear to be triggered by compensatory pathways and does not play a functional role (Supplemental Fig S4) may provide clues to the mechanisms underlying increased SHF addition in *lamb1a* mutants.

In the time frame in which heart morphogenesis and SHF addition occur, *lamb1b* is expressed throughout the endocardium at 30hpf (Fig S1B) and by 55hpf is restricted to the atrioventricular canal, the site of atrioventricular valve development (Fig 1J). This expression domain and dynamic is highly similar to other genes that are required for valvulogenesis such as *notch1b*, the expression of which is regulated by sensation of blood flow [24,63–66]. Furthermore, *fibronectin 1b*, a key ECM component required for valvulogenesis, is expressed in a similar domain to *lamb1b* at 55hpf and is also dependent on haemodynamic forces [34]. We therefore hypothesized that *lamb1b* expression is also flow-dependent, and that the misexpression of *lamb1b* throughout the endocardium of *lamb1a^Δ25^* mutants may reflect changes in cardiac function or flow sensitivity upon loss of laminin.

To investigate this, embryos from a *lamb1a^Δ25^* heterozygous incross were injected at the 1-cell stage with a translation-blocking morpholino oligonucleotide (MO) targeting *troponin T type 2a (cardiac)* (*tnnt2a*, previously *silent heart*) to block heart contractility and abolish blood flow [28]. Expression of *lamb1b* was then examined at 30hpf (Fig 5A-F), when uninjected and control *tp53* MO injected sibling embryos have low levels of endocardial *lamb1b* expression (Fig 5A,B), and uninjected and control injected *lamb1a^Δ25^* mutants misexpress *lamb1b* throughout the endocardium (Fig 5D,E) as previously described (Supplemental Fig 4). As expected, morpholino-mediated knockdown of *tnnt2a* in sibling embryos results in a loss of *lamb1b* expression in the endocardium (Fig 5C). Similarly, loss of heart contractility in *lamb1a^Δ25^* mutants results in reduced endocardial expression of *lamb1b* compared to uninjected and control mutants (Fig 5F). Together this demonstrates that endocardial expansion of *lamb1b* in *lamb1a^Δ25^* mutants is dependent on heart contractility.

**Figure 5.**
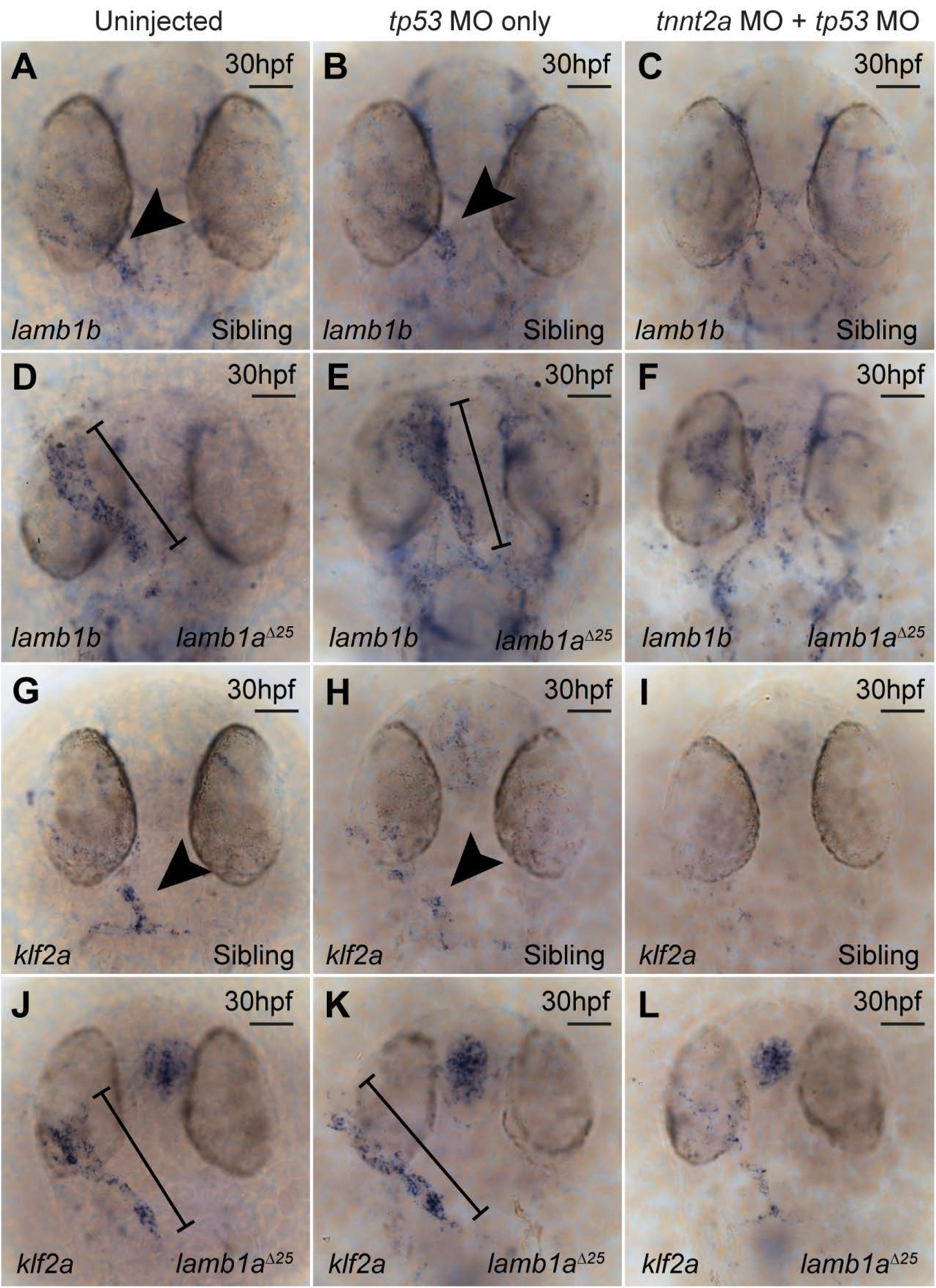
lamb1a mutants exhibit aberrant turbulent flow sensing. A-F: mRNA *in situ* hybridization analysis of *lamb1b* expression at 30hpf in sibling (A-C) and *lamb1a^Δ25^* mutant embryos (D-F), either uninjected (A,D), injected with a *tp53* MO (B,E) or *tp53* MO + *tnnt2a* MO (C,F). *lamb1b* is expressed predominantly in the ventricle/arterial pole of the heart tube endocardium in sibling uninjected (n=39/45) and control *tp53* MO-injected embryos (n=43/46) at 30hpf (arrowhead A,B), but is lost in embryos injected with *tnnt2a* MO (C, n=35/40). *lamb1b* expression is upregulated throughout the endocardium in *lamb1a^Δ25^* uninjected (n=23/23) and control *tp53* MO-injected mutant embryos (n=24/28) at 30hpf (bracket D,E) when compared with sibling controls (arrowhead A,B). Endocardial *lamb1b* expression is reduced in *lamb1a^Δ25^* mutant embryos injected with *tnnt2a* MO (F, n=24/28) when compared with control *lamb1a^Δ25^* mutants (D,E). G-L: mRNA *in situ* hybridization analysis of *klf2a* expression at 30hpf in sibling (G-I) and *lamb1a^Δ25^* mutant embryos (J-L), either uninjected (G,J), injected with a *tp53* MO (H,I) or *tp53* MO + *tnnt2a* MO (I,L). *klf2a* is expressed at low levels throughout the endocardium with elevated expression at the arterial pole in sibling uninjected (n=42/43) and control *tp53* MO-injected (n=37/39) embryos at 30hpf (arrowhead G,H), but is lost in embryos injected with *tnnt2a* MO (I, n=31/49). *klf2a* expression is upregulated particularly at the venous pole and atrium of *lamb1a^Δ25^* uninjected (n=23/25) and control *tp53* MO-injected mutant embryos (n=18/18) at 30hpf (bracket J,K) when compared with sibling controls (arrowhead G,H). Endocardial *klf2a* expression is reduced in *lamb1a^Δ25^* mutant embryos injected with *tnnt2a* MO (L, n=20/22) when compared with control *lamb1a^Δ25^* mutants (J,K). Dorsal views, anterior to top. Scale bars: 50μm.

To confirm altered flow-responsiveness in *lamb1a^Δ25^* mutant endocardium, we analysed the expression of *klf2a*, a key transcription factor whose expression is regulated by turbulent flow [24]. We injected embryos from a *lamb1a^Δ25^* heterozygous incross with *tnnt2a* MO and examined *klf2a* expression at 30hpf. In sibling uninjected and control embryos, *klf2a* expression is localised predominantly to the arterial pole endocardium (Fig 5G, H), whereas uninjected and control injected *lamb1a* mutants misexpress *klf2a* more broadly throughout the heart and also have prominent upregulation of *klf2a* in the head (Fig 5J,K). Morpholino-mediated knockdown of *tnnt2a* in sibling embryos results in almost total loss of *klf2a* expression in the endocardium compared to controls (Fig 5I). Similar to the effect on *lamb1b* expression, loss of heart contractility in *lamb1a^Δ25^* mutants results in reduced endocardial expression of *klf2a* compared to control mutants (Fig 5L). Together these data suggest that loss of Lamb1a may result in perturbations to the response to heart contractility and/or blood flow during heart looping morphogenesis.

The increase in SHF addition and altered expression of haemodynamic-responsive genes in *lamb1a^Δ25^* mutants suggest that cardiac function may play a role in the failure to restrict heart size in *lamb1a* mutants. We therefore investigated whether perturbing contractile and/or haemodynamic forces rescues cardiomegaly in *lamb1a^Δ25^* mutants by injecting the *tnnt2a* MO and measuring heart size (Fig 6A-D, Fig S6A & E). Blocking cardiac contractility in *lamb1a^Δ25^* mutant embryos significantly reduced heart size at 55hpf and 72hpf compared to control *lamb1a^Δ25^* mutants, suggesting that excess SHF addition is mediated by contractility upon loss of Lamb1a (Fig 6E, Fig S6A). To confirm this, we then examined the impact of loss of heart contractility specifically on SHF addition to the venous pole of the heart at 55hpf in *Tg(myl7:eGFP);Tg(myl7:DsRed)* transgenic embryos. In line with our previous data, uninjected and control-injected *lamb1a^Δ25^* mutant embryos display an increase in the number of newly-added SHF cells in the atrium at 55hpf (Fig 6K). However, in *lamb1a^Δ25^* mutants injected with *tnnt2a* MO, addition of SHF cells to the atrium is rescued to comparable levels with siblings (Fig 6K). Surprisingly, we also identified that loss of heart contractility in sibling embryos results in a subtle, yet significant increase in the number of GFP+/DsRed+ cells in the atrium at 55hpf (Figure 6J). This suggests that more broadly, heart contractility may be required to limit the rate of SHF addition to the atrium. Taken together, this demonstrates that Lamb1a is required to limit excessive, contractile-dependent SHF addition to the atrium during heart looping morphogenesis.

**Figure 6.**
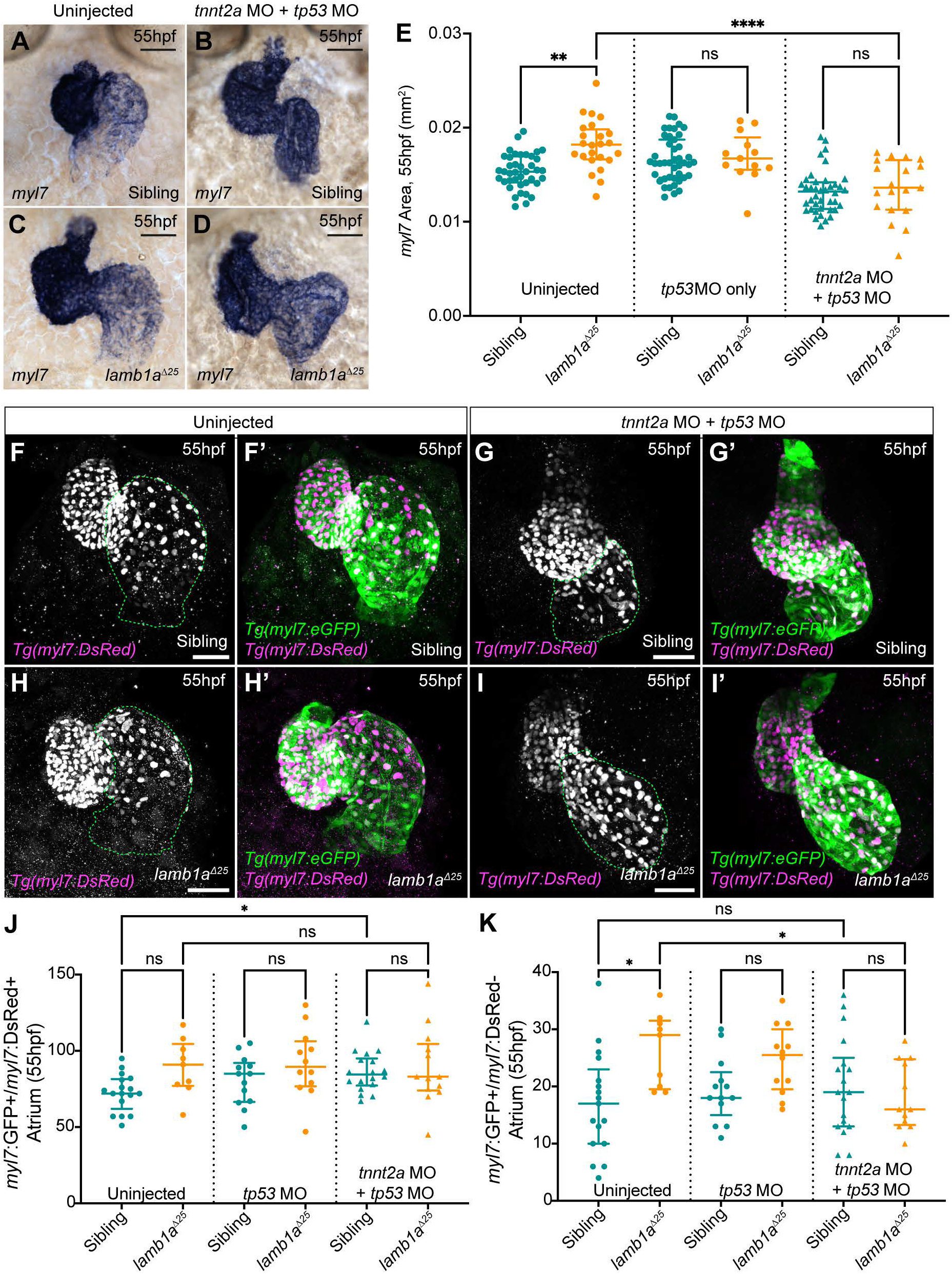
lamb1a limits excessive, contractility-dependent SHF addition to the venous pole. A-D: mRNA *in situ* hybridization analysis of *myl7* expression in sibling (A, B) and *lamb1a^Δ25^* mutant embryos (C,D) either uninjected (A,C) or injected with *tp53* MO *+ tnnt2a* MO (B,D). E: Quantification of *myl7* area in uninjected (sibling: n=40; *lamb1a^Δ25^*: n=24), *tp53* MO-injected control (sibling: n=43; *lamb1a^Δ25^*: n=13), and *tp53* MO + *tnnt2a* MO-injected (sibling: n=40; *lamb1a^Δ25^*: n=19) embryos at 55hpf. Horizontal bars indicate median with interquartile range, comparative statistics performed using Kruskal-Wallis test with multiple comparisons. F-I’: Maximum intensity projections of confocal image z-stacks in *Tg(myl7:eGFP);Tg(myl7:DsRed)* double transgenic sibling (F-G) and *lamb1a* mutant embryos (H-I) at 55hpf, either uninjected (F,H) or injected with *tp53* MO + *tnnt2a* MO (G,I). J-K: Quantification of double GFP+/DsRed+ cardiomyocytes (J) and GFP+/DsRed-SHF cells (K) at 55hpf in sibling and *lamb1a* mutant embryos either uninjected (sibling: n=17; *lamb1a^Δ25^*: n=9), injected with *tp53* MO (sibling: n=13; *lamb1a^Δ25^*: n=12), or injected with *tp53* MO + *tnnt2a* MO (sibling: n=18; *lamb1a^Δ25^*: n=13). Blocking heart contractility with the *tnnt2a* MO rescues excess SHF addition in *lamb1a* mutants (K). Scale bars: 50μm. Horizontal bars indicate median with interquartile range, comparative statistics performed using Brown-Forsythe and Welch ANOVA test with multiple comparisons, * = p < 0.05, ns = not significant.

Since heart contractility drives excessive SHF addition in *lamb1a^Δ25^* mutants, this could be due to increased heart rate. Importantly, analysis of heart rate at 2dpf and 3dpf did not reveal any significant differences between sibling and *lamb1a^Δ25^* mutant embryos (Supplemental Fig S6B), suggesting increased rate of heart contractility itself is not driving the aberrant contractility-mediated SHF addition. Additionally, since we observed changes in the expression of flow-responsive genes in *lamb1a^Δ25^* mutant embryos we wished to rule out that increased shear stress itself is responsible for cardiomegaly in *lamb1a* mutants. We lowered blood viscosity by injecting embryos from an incross of *lamb1a^Δ25^* heterozygous adults with a MO targeting the transcription factor *gata1a*, a master regulator of erythropoiesis [64,67]. Knockdown efficiency was confirmed by mRNA *in situ* hybridization expression analysis of the haemoglobin subunit *hbbe1.1* (*hemoglobin beta embryonic-1.1*) [68,69] in control and *gata1a*-injected embryos at 55hpf (Supplemental Fig S6C-E) and heart morphology was analysed at 72hpf by *myl7* expression. Quantification of heart area at 72hpf revealed that loss of Gata1a function and reduction in blood viscosity does not rescue *lamb1a^Δ25^* mutant heart size (Supplemental Fig S6F), demonstrating that cardiomegaly in *lamb1a* mutants is not due to increased flow sensing. Therefore, together our data demonstrates that loss of *lamb1a* results in excessive SHF addition to the atrium through a contractile-dependent, shear stress-independent mechanism.

### Retinoic Acid treatment during early SHF addition partially rescues cardiomegaly in lamb1a mutants

The molecular pathways underlying SHF patterning and addition are conserved among vertebrates, and include balanced, antagonistic, FGF8 and RA signalling across the arterial-venous axis of the cardiac-forming region [14]. Furthermore, cardiac function has been implicated in regulating the expression of *aldh1a2* (formerly *raldh2*), a key enzyme in the RA synthetic pathway [70], suggesting cross-talk between cardiac contractility and SHF patterning. We therefore examined the expression of *aldh1a2* (RA synthesis and RA signalling target) and *spry4* (FGF signalling responsive) in *lamb1a^Δ25^* mutants by mRNA *in situ* hybridization (Fig 7). At 30hpf and 55hpf, *lamb1a^Δ25^* mutants exhibit a marked upregulation of *aldh1a2* expression (Fig 7A-D) together with increased expression of *spry4* at 55hpf with variable penetrance (Fig 7E,F). As *aldh1a2* expression is sensitive to levels of RA in the embryo, where low levels of RA result in increased *aldh1a2* expression [71–73], this suggests that loss of Lamb1a alters the balance of FGF-RA signalling in the embryo during the window of SHF addition. To investigate whether the impact of loss of *lamb1a* on FGF and RA signalling is also contractility-dependent, we examined *aldh1a2* and *spry4* expression in sibling and *lamb1a^Δ25^* mutant embryos injected with *tnnt2a* MO. While uninjected and control injected *lamb1a^Δ25^* mutant embryos have a clear expansion of *aldh1a2* (Fig 7J,K), MO-mediated knockdown of *tnnt2a* abrogates endocardial *aldh1a2* expression in *lamb1a^Δ25^* mutants (Fig 7L). Surprisingly, blocking contractility in either sibling or *lamb1a* mutant embryos also resulted in altered *spry4* expression, with an expansion of *spry4* into the atrium that was most prominent upon loss of contractility in the *lamb1a^Δ25^* mutant (Fig 7M-R). Together this suggests, not only that disruptions to FGF-RA signalling in *lamb1a^Δ25^* mutants are partly regulated by heart function, but also that cardiac contractility itself can influence activity of the pathways regulating SHF addition.

**Figure 7.**
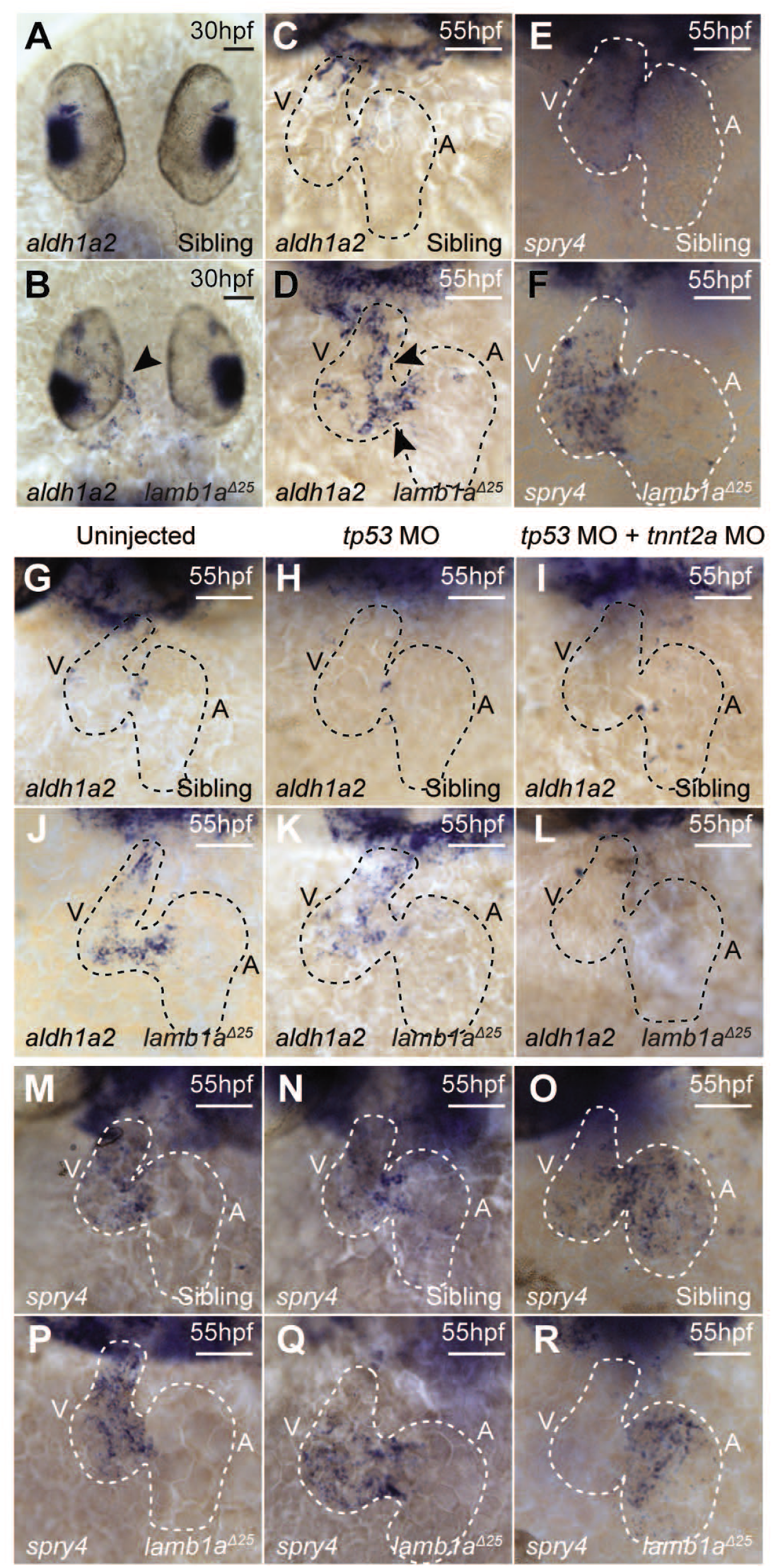
aldh1a2 upregulation lamb1a mutants is contractility-dependent. A-D: mRNA *in situ* hybridization analysis of *aldh1a2* expression in sibling and *lamb1a^Δ25^* mutant embryos at 30hpf (A,B) and 55hpf (C,D). *lamb1a^Δ25^* mutants exhibit an upregulation of *aldh1a2* expression in the endocardium at both stages (arrowheads B,D, 30hpf: n=14/20; 55hpf: n=16/17) when compared to siblings (30hpf: n=65/66; 55hpf: n=48/53). (E,F): mRNA *in situ* hybridization analysis of *spry4* expression at 55hpf reveals a mild upregulation of *spry4* expression in *lamb1a^Δ25^* mutants (F, n=10/21) when compared to sibling embryos (E, n=33/48). G-L: mRNA *in situ* hybridization analysis of *aldh1a2* expression at 55hpf in sibling and *lamb1a^Δ25^* mutant embryos, either uninjected (G,J), injected with *tp53* MO (H,K), or coinjected with *tp53* MO and *tnnt2a* MO (I,L). The upregulation of *aldh1a2* expression in the endocardium of *lamb1a^Δ25^* mutants (J: n=12/14, K: n=12/16) is lost upon injection with *tnnt2a* MO (L, n=12/18). M-R: mRNA *in situ* hybridization analysis of *spry4* expression at 55hpf in sibling and *lamb1a^Δ25^* mutant embryos, either uninjected (M,P), injected with *tp53* MO (N,Q), or coinjected with *tp53* MO and *tnnt2a* MO (O,R). *spry4* expression is expanded into the atrium in both sibling (n=20/30) and *lamb1a^Δ25^* mutant embryos (n=15/17) injected with *tnnt2a* MO (O,R), compared with ventricular expression in control embryos (M,N,P,Q). The mild upregulation of *spry4* expression in the ventricle of *lamb1a^Δ25^* mutants (P,Q) when compared to sibling embryos (M,N) is also lost in a subset of embryos upon injection of *tnnt2a* MO (R, n=9/17). A,B: Dorsal views, anterior to top. C-R: Ventral views, anterior to top. Black/white dashed line indicates heart outline. V - Ventricle, A - Atrium. Scale bars: 50μm.

Having established that excessive SHF addition in *lamb1a* mutants is dependent on heart contractility, and that RA signalling appears dysregulated upon loss of laminin in a contractility-dependent manner, finally we investigated whether we could rescue cardiomegaly in *lamb1a* mutants through modulation of RA. Studies in mouse have shown that loss of RA signalling in *aldh1a2* mutants results in hypoplastic atria [74,75], that upregulation of *aldh1a2* is a consequence of insufficient RA signalling [71–73], and that increased *spry4* expression suggests over-active FGF signalling (which is antagonised by RA signalling) [14]. Therefore, we hypothesized that addition of exogenous RA during early heart looping morphogenesis may rescue heart size in *lamb1a* mutants. We treated embryos from a *lamb1a^Δ25^* heterozygous incross from 24hpf to 55hpf with 100nM RA, washed the drug off and allowed the embryos to develop to 72hpf. We confirmed the efficacy of our drug treatment by *in situ* hybridization expression analysis of the RA-responsive gene *dhrs3a* [76–78], which is upregulated at 55hpf upon RA treatment (Supplemental Fig S7A-F). Analysis of heart size in sibling and *lamb1a^Δ25^* mutant embryos at 72hpf following 100nM RA treatment, revealed a significant partial rescue in cardiac size in *lamb1a* mutants (Supplemental Fig S7G-J,O) but a non-significant reduction in atrial area (Supplemental Fig S7K-N, P). This suggests that while contractility-dependent RA signalling in *lamb1a* mutants is perturbed, this is not the only pathway driving increased heart size, and that heart function likely affects SHF addition through additional mechanisms.

Together, we have shown the first requirement for laminins in regulating early vertebrate heart morphogenesis, promoting heart morphology and restricting heart size through restriction of SHF addition to the venous pole. Furthermore, our data indicate that the ECM and cardiac contractility together function to regulate the balance of SHF-related signalling pathways.

## Discussion

We have provided the first evidence that laminin restricts heart growth during looping morphogenesis by limiting the number of SHF cells incorporated into the venous pole of the heart. Previous studies have identified roles for ECM components such as Versican and Fibronectin in promoting SHF addition to the arterial pole of the heart [79–83], however we have identified an opposing role for Lamb1a in restricting excessive SHF addition to the venous pole. Highlighting the importance of cell-ECM interactions in the SHF loss of *Tbx1*, a master regulator of SHF addition, results in reduced expression of Integrin, loss of focal adhesion markers, and impaired filopodia formation in the SHF [84,85].

Heart function is tightly linked to heart morphology during development, and previous studies mainly focussed on the impact of contraction on regionalised ventricular cell shape change, valvulogenesis and trabeculation [31,63,86–88]. Our finding that excessive atrial SHF addition in *lamb1a* mutants can be rescued by abolishing heart contractility, but not shear stress (Fig 6, Fig S6), suggests that laminin may alter or dampen the physical force of heart contractility. *lamb1a* and *lamc1* are expressed broadly throughout the zebrafish embryo during the window of SHF migration, both within the heart and in the surrounding tissues where the SHF resides (Fig 1). Therefore whether the role of Lamb1a in restricting SHF addition to the atrium is autonomous or non-autonomous to the heart tube remains an open question.

The upregulation of flow-sensitive *klf2a* expression in *lamb1a* mutants suggests the dynamics of myocardial wall contraction may be altered. Since laminins coordinate ECM assembly, loss of laminin may alter ECM stiffness, which has been shown *in vitro* to impact cardiomyocyte contractility [89,90], while ECM composition can also affect the organisation of contractile apparatus in cardiomyocytes [91–93]. Alternatively, loss of laminin could affect how SHF cells interact with the underlying ECM, and studies in mouse have shown that the posterior SHF is under epithelial tension, which is important for cardiac growth [22]. This regionalised tension in the SHF is accompanied by nuclear localisation of the mechanosensitive transcription factor YAP [22], a phenomenon associated with increased ECM stiffness [94], and recent *in vitro* studies have demonstrated that laminin itself promotes nuclear YAP shuttling in keratinocytes [95]. Activation of YAP/TAZ signalling in the SHF at the venous pole of the heart is conserved in zebrafish [96], and the increased atrial SHF addition in 55hpf in *lamb1a* mutants is similar to that observed in *lats1/lats2* double mutants, which have a global increase in activity of YAP/TAZ signalling [96]. This suggests that ECM-mediated mechanotransduction in the SHF may represent a conserved mechanism regulating SHF addition that is disrupted in *lamb1a* mutants. How this is impacted by cardiac contraction is unclear, although it has been speculated that cardiac function could contribute to SHF tension [22], and it is conceivable that cardiomyocyte contractility contributes to the balance of pulling and pushing forces regulating SHF incorporation into the OFT in mouse [23]. Importantly, while blocking contractility rescues excessive cell addition to the venous pole in *lamb1a* mutants, morphology of the inflow tract appears impaired (Fig 6), suggesting that cardiac contraction is required to shape the venous pole.

In addition to facilitating cell-ECM interactions, laminins are crucial for ECM assembly, interacting with other ECM components such as Heparan Sulfate Proteoglycans (HSPGs). In turn, HSPGs interact both with additional ECM components such as Fibronectin (Fn), and with signalling molecules such as FGF in the extracellular space [38]. HSPG and Fn have both been previously shown to regulate FGF signalling during SHF addition to the OFT/arterial pole in mouse [97,98], therefore laminin may interact with HSPGs in the ECM to facilitate the signalling pathways regulating SHF addition. Supporting this we observed a mild upregulation of the FGF-response gene *spry4* in the ventricle of *lamb1a* mutant hearts at 55hpf (Fig 7). However, levels of FGF activity are also balanced by antagonistic RA signalling, the activity of which is regulated by the family of Cyp26 enzymes. This balance is crucial, as exemplified by the loss of ventricular SHF addition in Cyp26a1/Cyp26c1 deficient embryos, despite SHF progenitors being correctly specified and maintained [16]. We observe an upregulation of the RA-responsive RA-synthesising enzyme *aldh1a2* in *lamb1a* mutants as early as 30hpf (Fig 7), together suggesting that RA signalling is impaired upon loss of laminin, leading to a dysregulation of FGF activity. Providing some support for this hypothesis, timed RA treatments during early SHF addition partially rescued the increased heart size of *lamb1a* mutants at 3dpf (Supplemental Fig S7), although since the balance of RA-FGF is important, the global upregulation of RA is likely too broad to restore this balance completely. RA signalling during early development has recently been proposed to define the rate of cardiac progenitor differentiation in the anterior lateral plate mesoderm since disruption of RA signalling from 6hpf onwards results in a reduction in *ltbp3* expression and a loss of *isl1a*-positive pacemaker cells at the inflow tract [99]. Importantly, we do not observe changes in the size of *ltbp3* and *isl1a* expression domains at either the venous or arterial pole respectively at 30hpf (Fig S5), suggesting that altered FGF-RA signalling is not affecting the size of the SHF progenitor populations themselves and that RA signalling in *lamb1a* mutant hearts is disrupted after SHF specification. Furthermore, analysis of *aldh1a2* expression in *lamb1a* mutants in which cardiac contractility has been abrogated reveals that the *aldh1a2* upregulation in *lamb1a* mutants is dependent on heart function. Surprisingly, we also observe that loss of contractility disrupts the patterning of *spry4* expression, similar to a study demonstrating that muscle contractility regulates FGF signalling in the developing chick limb [100]. This suggests that dysregulation of these pathways may be secondary to altered contractility in *lamb1a* mutants, highlighting the complexity of interaction between the ECM, cardiac function, cell signalling, and SHF addition. Alternatively, other pathways regulating timely SHF differentiation may be altered in *lamb1a* mutants - for example *lamc1* promotes the correct localization of HSPGs, which in turn patterns BMP signalling during myotome development in zebrafish [101]. Together, a complex picture is emerging around the mechanisms underlying SHF deployment during heart development, and how the ECM regulates this process. While blocking contractility rescues excess SHF addition in *lamb1a* mutants, loss of contractility in a wild type embryo does not reduce cardiomyocyte or SHF number (Fig 6), suggesting that under wild type conditions heart function is not required for SHF addition to the atrium. However, we identify a mild but significant increase in the number of ‘older’ cardiomyocytes in the atrium of *tnnt2a* morphants, suggesting that contractility could play a role in the timing of SHF addition.

The tissue-restricted expression of individual laminin subunits we have identified during early heart morphogenesis suggests that different cell types in the heart may deposit specific laminin trimers which play distinct roles in heart morphogenesis and growth. We have identified multiple requirements for more broadly-expressed beta and gamma subunits during heart development, however our identification of tissue restricted alpha subunit expression (Figure 1 and S1) suggests that these alpha subunits may confer tissue-specificity to the multiple roles laminins play in the heart. Laminin deposition into the ECM by either myocardial or endocardial cell types likely facilitates signalling between these two tissue layers to coordinate cardiac growth. While the myocardium increases in cell number through SHF addition during heart looping morphogenesis [9,13], endocardial expansion is achieved through proliferation [32]. It has been proposed that endocardial growth is matched to myocardial chamber ballooning through Cadherin-5 mediated YAP-driven proliferation [102], and this is likely coordinated through the cardiac jelly. Detailed comparative analysis of cardiac growth, morphology, and function in *lama4* and *lama5* zebrafish mutants would therefore enable us to define how different cell types in the heart contribute to each process.

Both *LamB1* and *LAMB2* are detectable in human heart samples at gestational weeks 8/9, and are deposited into the ECM that surrounds the cardiomyocytes and the basement membrane of the endocardium [103]. Mutations in *lamc1* have been associated with Dandy-Walker Syndrome (DWS), a rare CNS disorder which is also associated with congenital heart defects [43]. We have shown a conserved requirement for zebrafish *lamc1* in heart morphogenesis with crispants also displaying hydrocephalus (Fig 2, Supplemental Fig S2), another symptom associated with DWS. Furthermore, our finding that loss of laminin may lead to altered contractility and heart morphology at relatively early stages of cardiac development may shed further light on the mechanisms underlying the progression of dilated cardiomyopathy in individuals with *LAMA4* mutations [41], since it is recognised that contractile dysfunction is a key factor in the initiation of cardiac remodelling and cardiomyopathies [104]. This conservation of laminin function highlights the value of zebrafish as a model for understanding the role of ECM dysfunction in human cardiac diseases. Moreover, the cardiac ECM is rapidly emerging as a key avenue for therapeutic targets following myocardial infarction (MI). In particular the zebrafish cardiac ECM is able to facilitate regeneration in human cardiac progenitors and in a mouse model of MI [105]. Changes in ECM composition in the mouse are associated with a loss of regenerative potential. Between P1 and P2, Collagen II, IV, elastin and laminin content dramatically increases in the heart, correlating with increased tissue stiffness and a dramatic decline in regenerative potential [106]. Our data demonstrating that *lamb1a* limits cardiomyocyte migration into the heart provides further avenues for investigation into how targeted modulation of the cardiac ECM can promote cardiomyocyte recruitment to the injury site, and improve regenerative potential.

Together, we describe the first direct evidence that laminins promote morphogenesis and growth during early vertebrate heart development, uncovering a novel role for laminin in restricting contractility-dependent SHF addition to the venous pole. This work also identifies new links between ECM composition, mechanical and biochemical cues in shaping the heart, reinforcing the importance of the extracellular environment during organ morphogenesis.

## Materials and Methods

### Zebrafish maintenance

Adult zebrafish were maintained according to standard laboratory conditions. The following, previously described lines were used: WT (AB), *Tg(myl7:eGFP)* [107], *Tg(myl7:lifeActGFP)* [108], *Tg(−5.1myl7:DsRed2-NLS)^f2^* [109], *grumpy^tj299a^* [56], *sleepy^sa379^* [110]. The lines generated for this study were: *lamb1a^Δ19^ (lamb1a^sh589^), lamb1a^Δ25^ (lamb1a^sh590^), lamb1b^promΔ183^ (lamb1b^sh587^), lamb1b^promΔ428^ (lamb1b^sh588^)*. Embryos were maintained in E3 medium at 28.5°C and were staged according to Kimmel et. al. [111]. Embryos older than 24hpf were transferred into E3 medium containing 0.003% 1-phenyl 2-thiourea (PTU, Sigma P7629) to inhibit pigment formation and aid imaging.

### CRISPR-Cas9-mediated lamc1 and lamb1a mutagenesis

*lamc1* and *lamb1a*-targeting gRNAs were designed using CHOPCHOP [112,113]. Following selection of suitable gene-specific sequence the first two nucleotides were converted from NG/GN to GG and the Protospacer Adjacent Motif (PAM) sequence (NGG) removed. The reverse complement of the resulting sequence was inserted into an ultramer scaffold sequence [114] containing a T7 promoter (AAAGCACCGACTCGGTGCCACTTTTTCAAGTTGATAACGGACTAGCCTTATTTTAACTTG CTATTTCTAGCTCTAAAACxxxxxxxxxxxxxxxxxxxxCTATAGTGAGTCGTATTACGC). The template was amplified by PCR (F: 5’-GCGTAATACGACTCACTATAG-3’, R: 5’-AAAGCACCGACTCGGTGCCAC-3’), and used as a template for *in vitro* transcription using MEGAshortscript T7 kit (Ambion/Thermo).

Generation of *lamc1* F0 crispants was carried out as previously described [55,115]. gRNAs were designed to target the initiation codon of *lamc1* (ENSDART00000004277.8) (*lamc1* F0 gRNA 1: 5’-GGCTTTCAATGCGACCGTGGTGG-3’ and *lamc1* F0 gRNA 2: 5’-GGCGTGCAGTCACGGAGCGATGG-3’). The injection mix containing gRNA, Cas9 protein and Phenol Red (Sigma P0290) was assembled on ice and incubated at 37°C for 5 minutes to aid Cas9-gRNA ribonucleoprotein complex formation for more efficient mutagenesis, prior to loading into a micro-injection needle. 1nL of Cas9-gRNA was injected into the yolk of 1-cell stage embryos, consisting of 500pg of each gRNA, 1.9nM Cas9 protein (NEB M0386T) and 14% Phenol Red. Efficacy of mutagenesis was confirmed through PCR amplification of the targeted region of *lamc1* genomic DNA (F: 5-ATCAAGACAGTGACGGTAGCAA-3’, R: 5’-TGTGGCATGATTTAGTGACTCC-3’) and uninjected, gRNA-only injected and Cas9-only injected embryos were included as controls.

To generate *lamb1a* (ENSDART00000170673.2) mutant zebrafish, a single gRNA targeting Exon 6 (5’-GGATCCTCAATCCTGAAGGCAGG-3’) was injected together with Cas9 protein and Phenol Red into the yolk at the 1-cell stage. Each embryo was injected with 2ng gRNA, 1.9nM Cas9 protein and 10% Phenol Red. CRISPR Cas9-injected embryos were raised to adulthood (F0) and outcrossed to wildtype to identify F0 individuals with germline transmission of deletions that result in a frameshift and subsequent premature termination codon. Embryos from F0 outcrosses were genotyped by PCR to amplify the region of *lamb1a* targeted for mutagenesis (F: 5’-CTTCTGTCTCTCATGGGCCA-3’, R: 5’-TGCCTTTACTTTGAATTCTGGGG-3’), and mutations analysed by Sanger sequencing. Two *lamb1a* coding sequence deletion alleles were recovered: *lamb1a^Δ19^* (*lamb1a^sh589^*) and *lamb1a^Δ25^* (*lamb1a^sh590^*). F0 founders transmitting these mutations were outcrossed to WT (AB) and offspring raised to adulthood. Phenotypic analyses were carried out using F2 or F3 adults.

### Morpholino-mediated knockdown

All morpholinos used are previously described: *tp53-MO* [116], *tnnt2a-MO* [28], *gata1a-MO* [117]. *tp53* and *tnnt2a* morpholinos were purchased from GeneTools and resuspended in MilliQ to 1mM. The following concentrations were used for knockdown: *tp53* 250nM and *tnnt2a* 125nM. The *gata1a* morpholino was a gift from J. Serbanovic-Canic, and injected at 200nM. *tnnt2a* or *gata1a* morpholinos were co-injected with the *tp53* morpholino. Embryos were injected with 1nL of morpholino solution into the yolk at the 1-cell stage.

### mRNA in situ hybridization

For chromogenic mRNA *in situ* hybridization embryos were fixed overnight in 4% paraformaldehyde (PFA, Cell Signalling Technology #12606), for fluorescent mRNA *in situ* hybridization embryos were fixed overnight in 4% paraformaldehyde containing 4% sucrose. Following fixation embryos were washed 3 x 5 min in PBST and transferred into 100% MeOH for storage at −20°C. Chromogenic mRNA *in situ* hybridization was carried out as previously described [118]. Fluorescent *in situ* hybridizations were carried out using the Perkin-Elmer TSA kit [119]. The Fluo-labelled *fli1* riboprobe was developed with Tyr-Cy5, followed by the Dig-labelled gene of interest riboprobe with Tyr-Cy3. Following probe signal amplification, embryos were fixed in 4% PFA with sucrose overnight then washed into PBST for immunohistochemistry. The following previously published probes were used: *lamb1b* [53], *fli1* [120], *myl7* [121], *myh6* [52], *myh7l* [52], *aldh1a2* [122], *spry4* [123], *ltbp3* [10], *klf2a* [124] and *hbbe1.1* [69]. All other probes were generated for this study - see supplementary methods for details. Riboprobes were transcribed from a linearized template in the presence of DIG-11-UTP or Fluorescein-11-UTP (Roche).

### Immunohistochemistry

Embryos were fixed overnight in 4% paraformaldehyde containing 4% sucrose, washed 3 x 5 mins in PBST and transferred into 100% MeOH for storage at −20°C. Embryos were rehydrated into PBST, washed briefly in PBST and 2 x 5 mins in PBS-Triton (0.2% Triton-X in PBS). Embryos were incubated in blocking buffer (10% Goat Serum (Invitrogen) in PBS-Triton) at room temperature with gentle agitation for 1 hour. Blocking buffer was removed and replaced with blocking buffer containing 1% DMSO and primary antibodies. Embryos were incubated overnight at 4°C with gentle agitation. Following removal of primary antibodies, embryos were extensively washed in PBS-Triton and incubated in blocking buffer containing 1% DMSO and secondary antibodies overnight at 4°C with gentle agitation. After removal of secondary antibodies, embryos were extensively washed in PBS-Triton at room temperature before being prepared for imaging. The following antibodies were used: Chicken anti-GFP (1:500, Aves lab) and Rabbit anti-DsRed (1:200, Takara 632496), Donkey anti-Chicken-Cy2 (1:200, Jackson labs) and Goat anti-Rabbit-Cy3 (1:200, Jackson labs).

### Retinoic Acid treatments

Retinoic Acid powder (R2625-50MG) was dissolved in DMSO (Sigma 276855) to a stock concentration of 10mM, and aliquots stored at −80C. Embryos were manually dechorionated prior to treatment, and 10 *lamb1a* mutant and 13 *lamb1a* sibling embryos were placed in a glass petri dish. Stock RA was diluted 1:10,000 in E3-PTU to give a working concentration of 100nM RA with 1% DMSO, and 8mL was added to treatment dishes. Control embryos were incubated with either E3-PTU or E3-PTU with 1% DMSO. Embryos were incubated in RA or control medium from 24hpf to 55hpf, when the drug was removed by rinsing embryos 3 x 5 min in E3, and either fixed immediately or development allowed to proceed until 72hpf and then fixed. Embryos were protected from light during the treatment window. Each RA treatment/control treatment was treated as one experimental unit for quantification, with an average value calculated from all embryos for each treatment. These treatment averages then formed one experimental replicate for statistical analyses, and treatments were replicated four times.

### Imaging and image quantification

Prior to quantification files were blinded using an ImageJ Blind_Analysis plugin (modified from the Shuffler macro, v1.0 26/06/08, Christophe LeTerrier). Looping ratio, heart area and chamber area were quantified as previously described [52].

Total heart cardiomyocyte cell number and internuclear distance was quantified from *Tg(myl7:DsRed)* transgenic hearts. Z-stacks of fixed hearts were imaged on a Nikon A1 confocal, using a 40x objective with a z-resolution of 1μM. The DsRed channel of each heart was used to generate a depth-coded z-projection of the z-stack, using the temporal colour code function in Fiji. Cell number in the atrium and ventricle were quantified from these z-projections. Internuclear distance was quantified by measuring the distance between DsRed+ nuclei with the same or similar depth-coding in the projection. Six cells were selected per chamber, and from each cell the distance to the four nearest neighbours with similar z positions was measured. Average internuclear distance was then calculated for each chamber in each embryo.

Atrial second heart field addition was quantified similar to previous methods [13]. Stacks were opened in Fiji and converted to Maximum Intensity Projections. Using the DsRed channel only, the intensity was increased to maximum and the number of atrial DsRed+ nuclei were quantified using the ROI Manager, the GFP channel was used to confirm position in the heart. Using the GFP channel only, the intensity was increased to maximum and GFP+ nuclei not previously counted in the ROI Manager were quantified as DsRed-. Returning to the original stack, individual slices were examined, together with the ROIs for DsRed+ and DsRed-atrial cells to ensure that no cells had been missed or miscounted. Cells derived from the ventricle or atrioventricular canal were discounted.

## Acknowledgements

We thank Tanya Whitfield and Juliana Sánchez-Posada for helpful comments on the manuscript. Additional imaging work was performed at the Wolfson Light Microscopy Facility using a Nikon A1 microscope. C.J.D would like to thank Martin and Jane Derrick for financial support during 2020.

