## Supplemental Material for "Lamb1a regulates atrial growth by limiting excessive, contractility-dependent second heart field addition during zebrafish heart development"

**Supplemental Data**

Containing:

Supplemental Figures S1 - S7

Supplemental Methods

**Supplemental Figures**

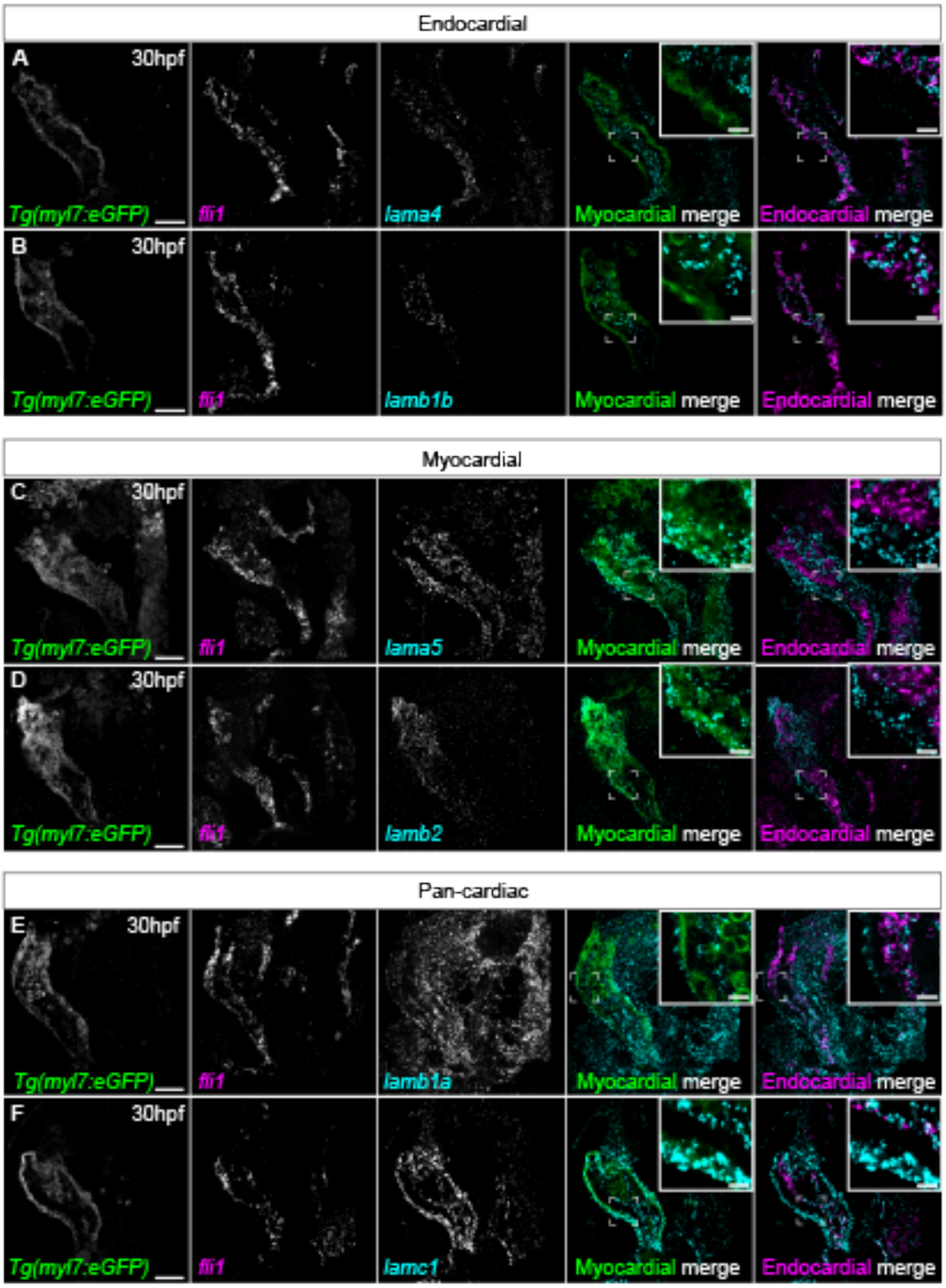


*Supplemental Fig S1 - Laminin subunits exhibit tissue-specific expression*

A-F: Single z-plane confocal images of fluorescent mRNA *in situ* hybridization analysis of *lama4*, *lamb1b, lama5, lamb2, lamb1a* or *lamc1* expression (cyan) in *Tg(myl7:eGFP)* embryos (myocardium, green) counterstained for *fli1* mRNA (endocardium, magenta). *lama4* and *lamb1b* expression colocalises with *fli1* in the endocardium (A,B), while *lama5* and *lamb1a* are expressed in the myocardium (C,D). *lamb1a* and *lamc1* are expressed in both myocardial and endocardial cells (E,F). Scale bars main panels: 50µm, insets: 10µm.

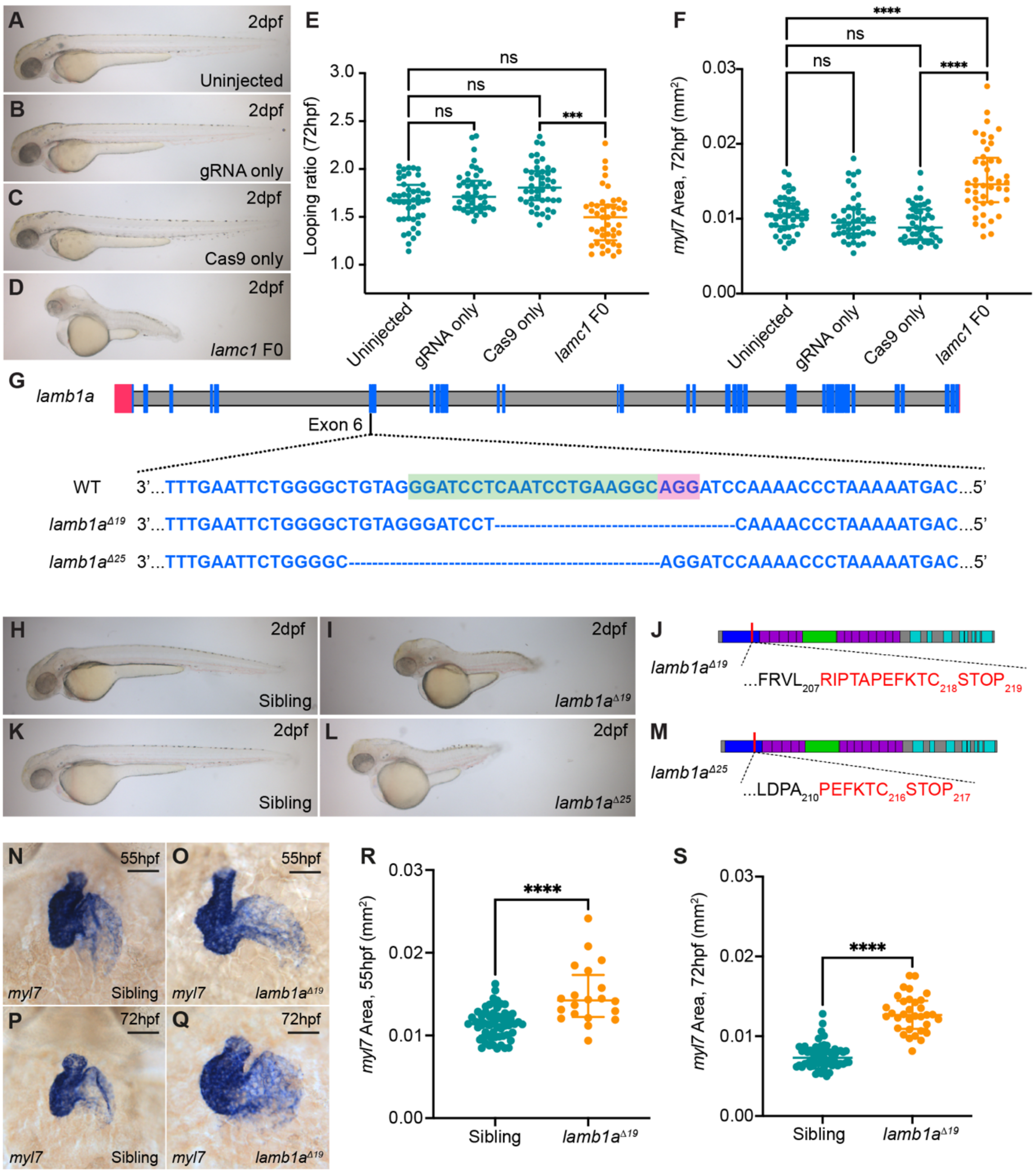


*Supplemental Fig S2 - lamc1 and lamb1a mutants have defects in heart morphogenesis*

A-D: Brightfield images of uninjected control (A), gRNA- or Cas9-only injected controls (B,C) or *lamc1* F0 crispant embryos (D) at 2dpf. Lateral views, anterior to left. E-F: Quantification of looping ratio (E) and heart size (F) in uninjected embryos (n=47), gRNA only control embryos (n=44), Cas9 only control embryos (n=44), and *lamc1* crispants (n=44) at 72hpf. Kruskal-Wallis test. G: Schematic depicting *lamb1a* gene (danRer10/GRCz10), with non-coding exons in red and coding exons in blue. gRNA target site (green) is located in exon 6 of wild type (WT) *lamb1a* (magenta indicates PAM sequence), and 2 deletion alleles of 19bp and 25bp were recovered. H-L: Brightfield images of sibling embryos (H,K), *lamb1a*$^{\Delta 19}$ (I) and *lamba*$^{\Delta 25}$ mutant embryos (L) and 2dpf. J,M: Schematic representation of Lamb1a protein structure (UniProt Q8JHV7), with the alterations in amino acid sequence depicted in red. Both *lamb1a* mutant alleles result in frameshift and insertion of a stop codon.

N-Q: mRNA *in situ* hybridization analysis of *myl7* expression in sibling (N,P) and *lamb1a*$^{\Delta 19}$ mutant embryos (O,Q) and 55hpf and 72hpf. Ventral views, anterior to top. R-S: Quantification of *myl7* area reveals a significant increase in heart size in *lamb1a* mutants (55hpf: n=20; 72hpf: n=33) when compared to siblings (55hpf: n=62; 72hpf: n=62) at 55hpf and 72hpf. Mann-Whitney test. **** = $p < 0.0001$, *** = $p < 0.001$, ** = $p < 0.01$. Scale bars: 50µm.

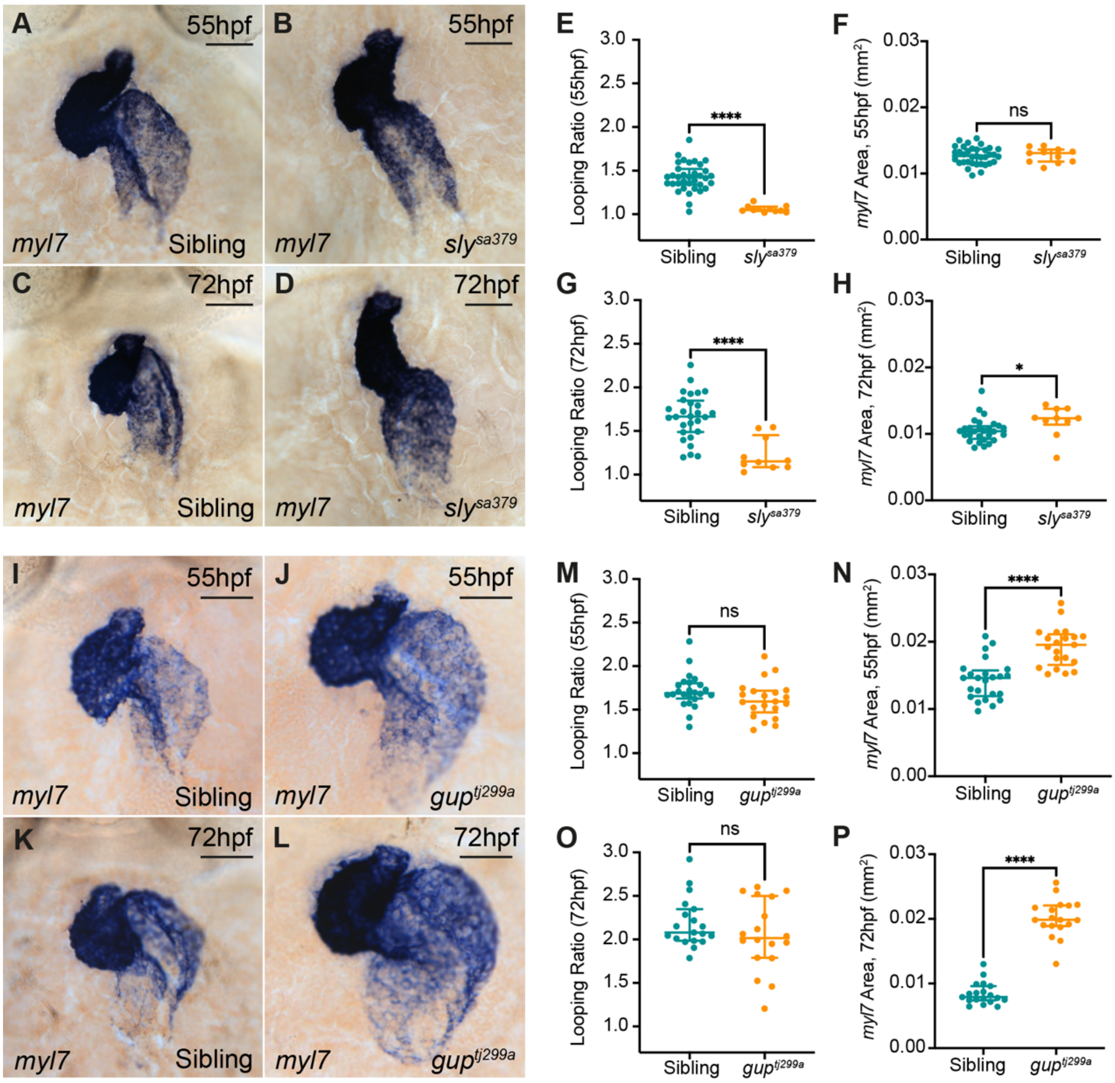


*Supplemental Fig S3 - sleepy and grumpy mutants recapitulate lamc1 crispant and lamb1a mutant heart phenotypes*

A-D: mRNA *in situ* hybridization analysis of *myl7* expression in sibling (A: n=35; C: n=28) and *sly*$^{sa379}$ mutant embryos (B: n=11; D: n=10) at 55hpf and 72hpf. Ventral views, anterior to top. E-H: Quantification of looping ratio in sibling and *sly*$^{sa379}$ mutant embryos reveals a reduction in heart looping morphology at both 55hpf (E) and 72hpf (G). Heart size is unaffected in *sly*$^{sa379}$ mutant embryos compared to siblings at 55hpf (F), however *sly*$^{sa379}$ mutants have slightly enlarged hearts at 72hpf (H). I-L: mRNA *in situ* hybridization analysis of *myl7* expression in sibling (I: n=24; K: n=19) and *gup*$^{tj299a}$ mutant embryos (J: n=21; L: n=18) at 55hpf and 72hpf. Ventral views, anterior to top. M-P: Quantification of looping ratio reveals no significant difference in looping morphology between sibling and *gup*$^{tj299a}$ mutant embryos (M,O). However, *gup*$^{tj299a}$ mutant embryos exhibit enlarged hearts when compared to siblings at both 55hpf (N), and 72hpf (P). All statistical analyses performed using Mann-Whitney test, **** = $p < 0.0001$, *** = $p < 0.001$, ** = $p < 0.01$, * = $p < 0.05$, ns = not significant. Scale bars: 50µm.

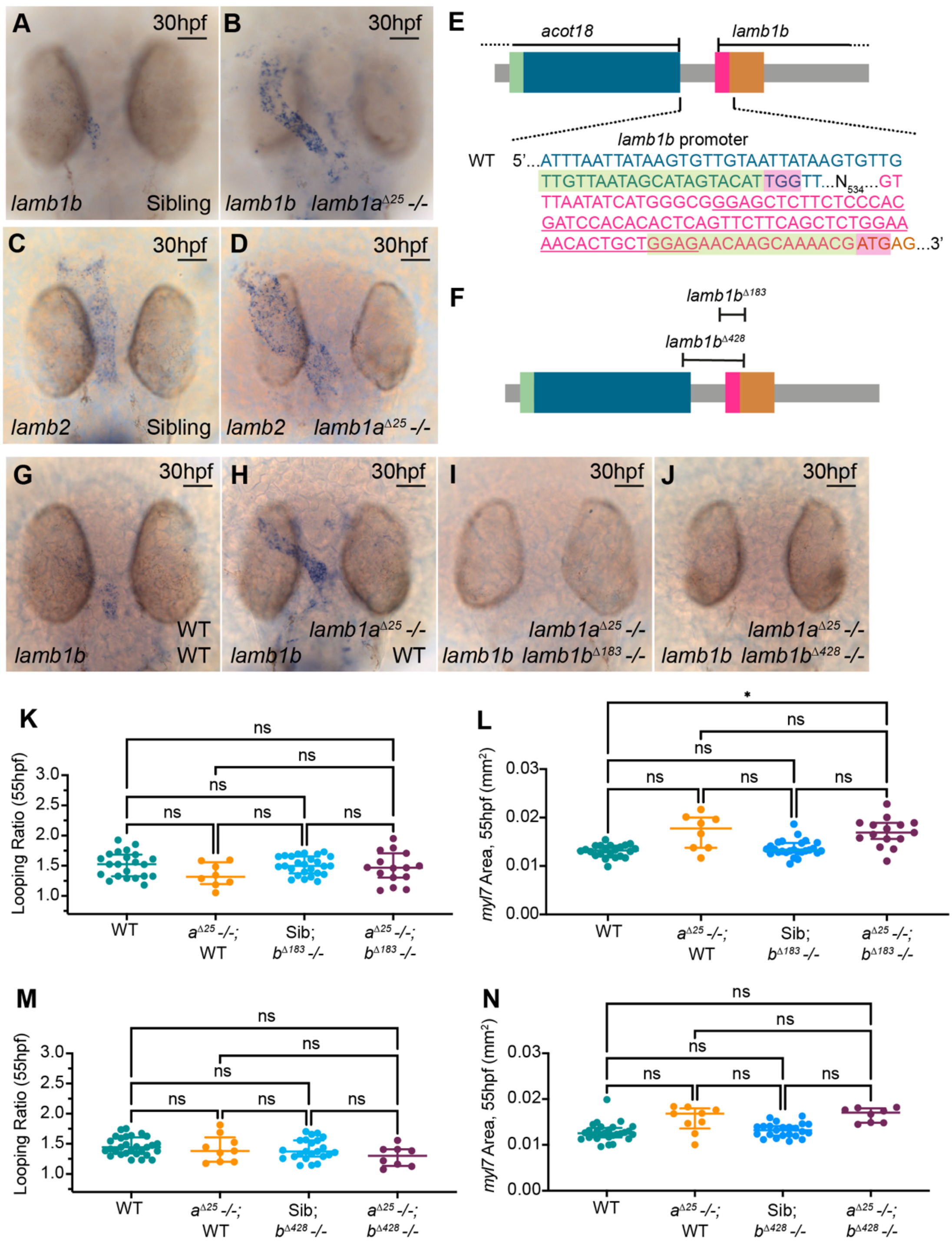


*Supplemental Fig S4 - lamb1b is dispensable for heart development*

A-D: mRNA *in situ* hybridization expression analysis of *lamb1b* (A,B) and *lamb2* (C,D) expression in sibling (A,C) and *lamb1a*$^{\Delta 25}$ mutant embryos (B,D). *lamb1a* mutants exhibit an upregulation of *lamb1b* throughout the endocardium (B, n=15/16) compared to wild types (A, n=15/17), but *lamb2* expression in the myocardium is unaffected in *lamb1a*$^{\Delta 25}$ mutants (D, n= 16/20) when compared to siblings (C, n=10/13) Dorsal views anterior to top. E-F: Schematic representation of CRISPR gRNAs (green sequence, PAM in magenta) targeting the promoter of *lamb1b* (danRer10/GRCz10). Two *lamb1b* deletion alleles were recovered: *lamb1b*$^{\Delta 183}$ and *lamb1b*$^{\Delta 428}$. G-J: mRNA *in situ* hybridization analysis of *lamb1b* expression in wild type (G), *lamb1a*$^{\Delta25}$ mutants (H), *lamb1b*$^{\Delta183}$*;lamb1a*$^{\Delta25}$ double mutants (I), or *lamb1b*$^{\Delta428}$*;lamb1a*$^{\Delta25}$ double mutants (J). *lamb1b* expression is abrogated in both *lamb1b*$^{\Delta183}$*;lamb1a*$^{\Delta25}$ (n=6/6) and *lamb1b*$^{\Delta428}$*;lamb1a*$^{\Delta25}$ double mutants (n=6/6) (I,J). Dorsal views, anterior to top. K-L: Quantification of heart looping ratio (K) and *myl7* expression domain as a proxy for heart size (L) in wild type (n=22), *lamb1a*$^{\Delta25}$ single mutants (n=8), *lamb1b*$^{\Delta183}$ single mutants (n=26), and *lamb1b*$^{\Delta183}$*;lamb1a*$^{\Delta25}$ double mutant embryos (n=15) at 55hpf. M-N: Quantification of heart looping ratio (M) and heart size (N) in wild type (n=25), *lamb1a*$^{\Delta25}$ single mutants (n=9), *lamb1b*$^{\Delta428}$ single mutants (n=23), and *lamb1b*$^{\Delta428}$*;lamb1a*$^{\Delta25}$ double mutant embryos (n=8) at 55hpf. Loss of *lamb1b* in *lamb1a*$^{\Delta25}$ neither induces defects in heart looping morphology, nor rescues heart size, in *lamb1a*$^{\Delta25}$ mutants. All statistical analyses performed using Kruskal Wallis test, * = $p < 0.05$, ns = not significant. Scale bars: 50µm.

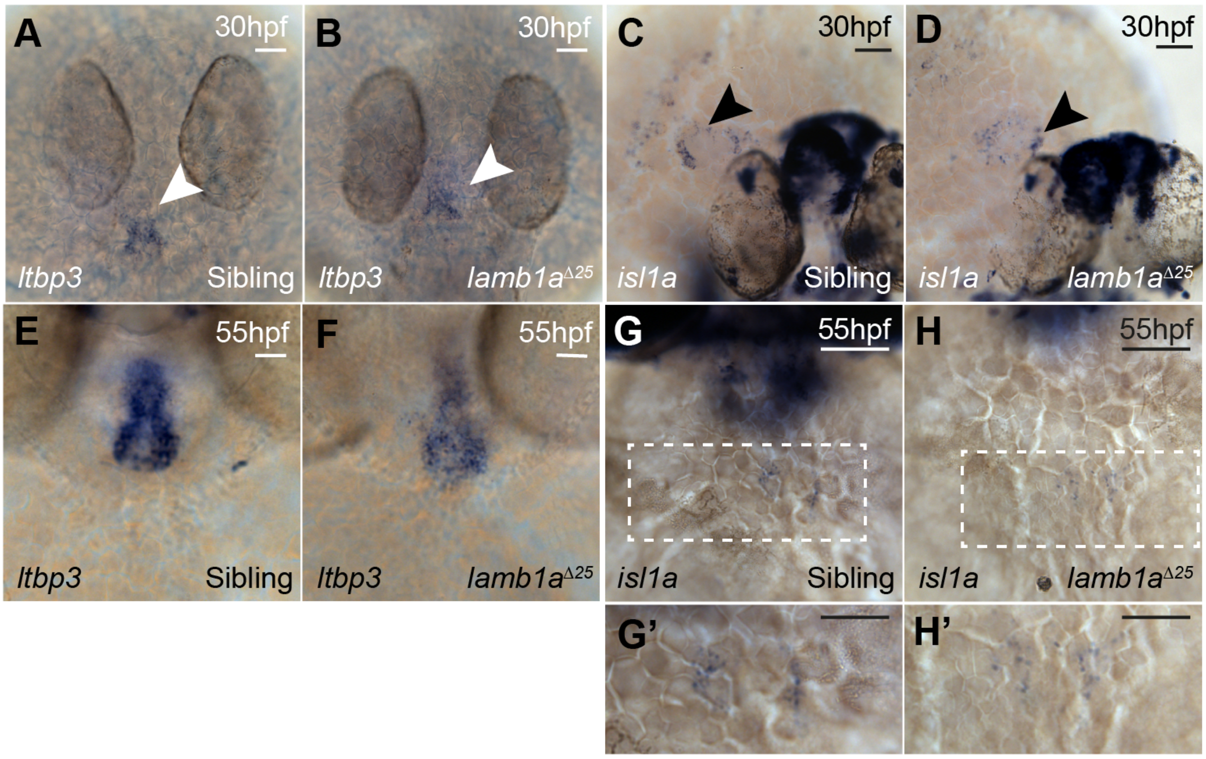


*Supplemental Fig S5 - SHF markers are not expanded in lamb1a mutant embryos*

A,B,E,F: mRNA *in situ* hybridisation analysis of *ltbp3* expression at 30hpf (A-B) and 55hpf (E-F) in sibling and *lamb1a*$^{Δ25}$ mutant embryos. The domain of *ltbp3* expression in the SHF at the arterial pole of the heart (white arrowhead) is unaffected in *lamb1a* mutants (B) when compared with siblings (A) at 30hpf. At 55hpf the size of the *ltbp3* expression domain is unaffected in *lamb1a* mutants (F, n=23/26), although staining levels are slightly reduced when compared with sibling embryos (E, n=35/46). C,D,H,I: mRNA *in situ* hybridization analysis of *isl1a* expression at 30hpf (C,D) and 55hpf (G,H) in sibling and *lamb1a*$^{Δ25}$ mutant embryos. The domain of *isl1a* expression at the venous pole of the heart (black arrowhead) is unaffected in *lamb1a*$^{Δ25}$ mutants (D, n=19/23; H, n=17/23) when compared to sibling embryos (C, n=43/54; G, n=38/47) at both stages. G',H': Higher magnification of venous pole region contained in white box in G and H.

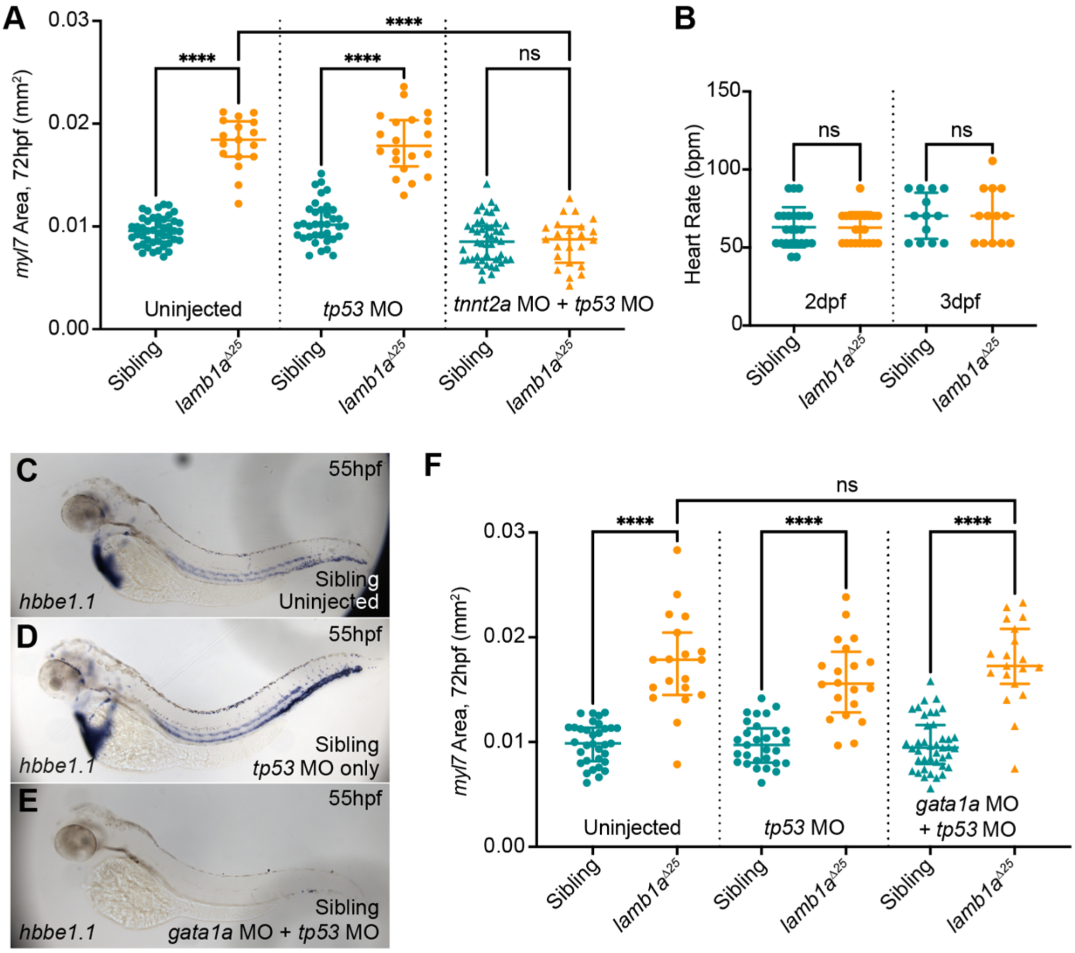


*Supplemental Fig S6 - lamb1a restricts cardiac growth independent of haemodynamic force*

A: Quantification of *myl7* expression domain at 72hpf as a proxy for heart size in sibling and *lamb1a*$^{\Delta25}$ mutants, either uninjected (sibling: n=45; *lamb1a*$^{\Delta25}$ mutants: n=17), injected with *tp53* MO (sibling: n=36; *lamb1a*$^{\Delta25}$ mutant: n=20) or co-injected with *tp53* MO and *tnnt2a* MO (sibling: n=46; *lamb1a*$^{\Delta25}$ mutant: n=23). Cardiomegaly is rescued in *lamb1a*$^{\Delta25}$ mutants injected with *tnnt2a* MO. B: Quantification of heart rate in sibling and *lamb1a*$^{\Delta25}$ mutant embryos at 2dpf and 3dpf reveals that *lamb1a*$^{\Delta25}$ mutants do not exhibit an elevated heart rate. C-D: mRNA *in situ* hybridization analysis of *hbbe1.1* expression at 55hpf in sibling embryos, either uninjected (C), *tp53* MO (D), or *tp53* MO and *gata1a* MO (E). Injection of *gata1a* MO prevents the formation of erythroid cells (E). Lateral views, anterior to left. F: Quantification of *myl7* expression domain as a proxy for heart size in sibling and *lamb1a*$^{\Delta25}$ mutants, either uninjected (sibling: n=33; *lamb1a*$^{\Delta25}$ mutants: n=19), injected with *tp53* MO (sibling: n=29; *lamb1a*$^{\Delta25}$ mutant: n=20) or co-injected with *tp53* MO and *gata1a* MO (sibling: n=41; *lamb1a*$^{\Delta25}$ mutant: n=19) reveals that loss of erythroid cells through injection of *gata1a* MO does not rescue heart size in *lamb1a*$^{\Delta25}$ mutant embryos. All statistical analyses performed using Mann-Whitney test, **** = $p < 0.0001$, ns = not significant. Scale bars: 50µm.

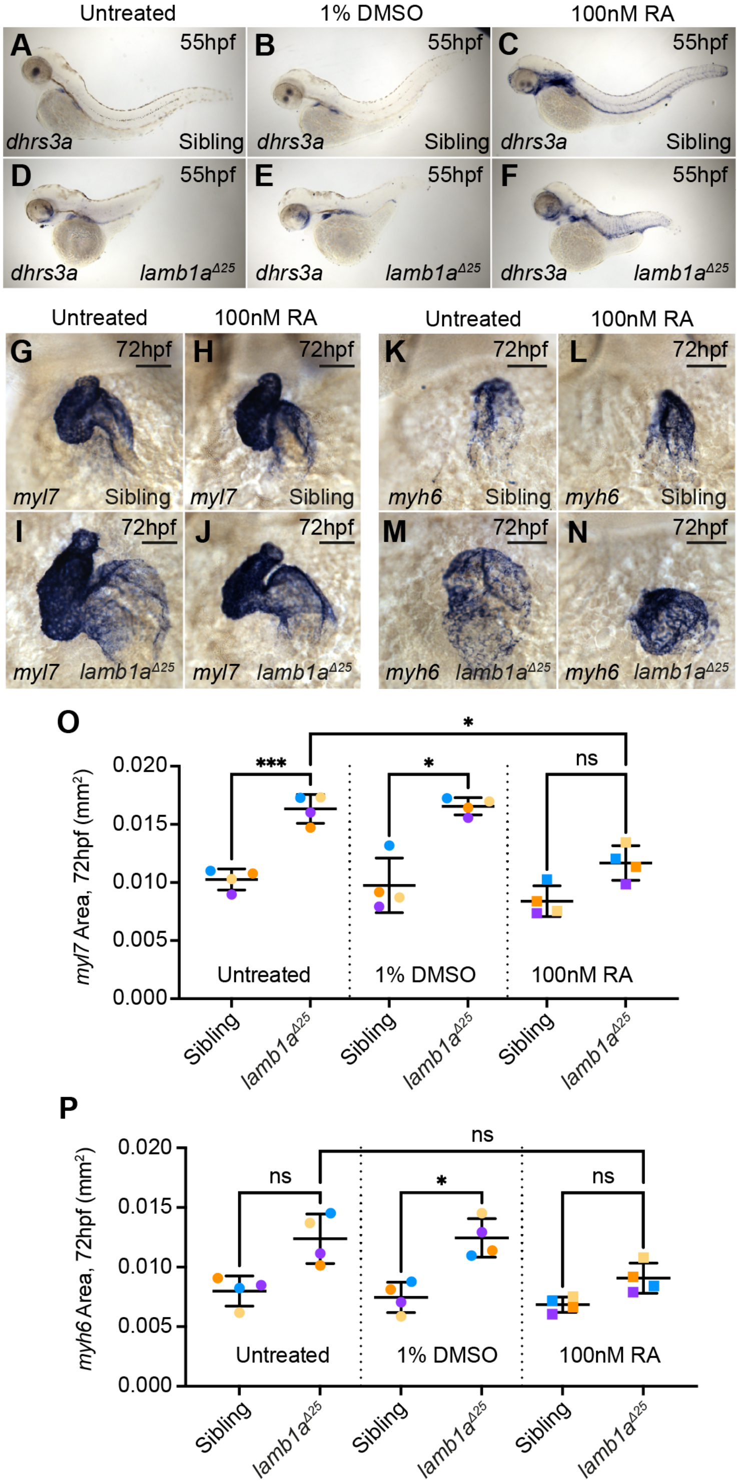


*Supplemental Fig S7 - Retinoic Acid treatment during SHF addition partially rescues cardiomegaly in lamb1a mutant embryos*

A-F: mRNA *in situ* hybridization analysis of *dhrs3a* expression at 55hpf in sibling and $lamb1a^{\Delta25}$ mutants either untreated, incubated with DMSO, or incubated with 100nM RA from 24hpf to 55hpf. RA treatment results in an upregulation of the RA-responsive gene *dhrs3a* (C, F) compared to untreated controls (A,B,D,E). Lateral views, anterior to left). G-J: mRNA *in situ* hybridization expression analysis of *myl7* expression at 72hpf in sibling and $lamb1a^{\Delta25}$ mutant embryos, either untreated (G,I) or incubated in 100nM RA between 24hpf and 55hpf (H,J). K,N: mRNA *in situ* hybridization expression analysis of *myh6* at 72hpf in the atrium of sibling and *lamb1a*$^{\Delta 25}$ mutant embryos, either untreated (K,M) or incubated in 100nM RA between 24hpf and 55hpf (L,N). Ventral views, anterior to top. O-P: Quantification of *myl7* expression domain (O) and *myh6* expression domain (P) in control and RA-treated embryos. RA treatment significantly reduces heart size in *lamb1a* mutants compared to controls (O). All statistical analyses performed using Kruskal Wallis test, *** = $p < 0.001$, * = $p < 0.05$, ns = not significant. Scale bars: 50µm.

**Supplemental Methods**

*mRNA in situ hybridization probe generation.*

Probe constructs were generated through PCR amplification of a DNA fragment from total zebrafish cDNA at 55hpf, and ligated into either the pCRII-TOPO vector or pCR4-TOPO vector (ThermoFisher).

| Probe | Primer Sequence | Accession Number; ZFIN ID) |
|---|---|---|
| *lama4* | F: 5’-CGATCAACTGCAGAGACACG-3’ | ENSDARG00000020785; ZDB-GENE-040724-213 |
| | R: 5’-GATGAACTTCTGCTCGGCTG-3’ | |
| *lama5* | F: 5’-CCCTCGCACCAATACATGTG-3’ | ENSDARG00000058543; ZDB-GENE-030131-9823 |
| | R: 5’-CATTGGGTCTGCATCGACAG-3’ | |
| *lamb1a* | F: 5’-TCCACTTCACCCACCTCATC-3’ | ENSDARG00000101209; ZDB-GENE-021226-1 |
| | R: 5’-GGTCACAGTTCCTTCCGGTA-3’ | |
| *lamb2* | F: 5’-CAAGACAACCGAAGCCAACA-3’ | ENSDARG00000002084; ZDB-GENE-081030-4 |
| | R: 5’-GGCTTACAGTCAGGGAAGGT-3’ | |
| *lamc1* | F: 5’-GTGCTCTTGTAATCCAGCCG-3’ | ENSDARG00000036279; ZDB-GENE-021226-3 |
| | R: 5’-GCTCACATCGCTTACCTGTG-3’ | |
| *isl1a* | F: 5’-GGACCTAACACCGCCTTACT-3’ | ENSDARG00000004023; ZDB-GENE-980526-112 |
| | R: 5’-TAGGACTCGCTACCATGCTG-3’ | |
| *dhrs3a* | F: 5’-AAAGGTGATTTTGTGGGGCC-3’ | ENSDARG00000044982; ZDB-GENE-040801-217 |
| | R: 5’-AACAAGCCATCTCGATTCGC-3’ | |

*Generation of lamb1b promoter mutants*

Two gRNAs were designed to target the upstream of the annotated promoter of *lamb1b* (ENSDARG00000045524) according to the Eukaryotic Promoter Database (Dreos 2014 and Dreos 2017) (*lamb1b* crRNA 1: 5’-TTGTTAATAGCATAGTACATTGG-3’ underlining denotes PAM) and downstream of the annotated initiation codon (*lamb1b* crRNA 2: 5’-GGAGAACAAGCAAAACGATGAGG-3’ underlining denotes PAM). Sequence-specific CRISPR RNAs (crRNA) were synthesised by Merck and resuspended in MilliQ water to 500uM and dilutions made for working stocks. 2nL of a Cas9-gRNA Ribonucleoprotein complex was then injected into the yolk of 1-cell stage embryos. Each embryo was injected with 61.2nM of crRNA, 122.5nM tracrRNA, 3.9nM Cas9 and 14% Phenol Red.

CRISPR Cas9-injected embryos were raised to adulthood (F0) and individual adults were outcrossed to wildtype to identify germline transmission of suitable promoter deletions. Embryos collected from these outcrosses were genotyped by PCR to amplify the region of *lamb1b* targeted for mutagenesis (Forward: 5'-TCACACTAAGACATGGGGCA-3', Reverse: 5'-ACCAAGCAACCAAAACACTGA-3'). Successful promoter deletion was identified by presence of a smaller PCR fragment by gel electrophoresis and subsequent Sanger sequencing of the PCR fragment to confirm the deletion. Two separate *lamb1b* promoter deletion alleles were recovered: $lamb1b^{prom\Delta183}$ ($lamb1b^{sh587}$) and $lamb1b^{prom\Delta428}$ ($lamb1b^{sh588}$). F0 founders transmitting these mutations were outcrossed to $lamb1a^{\Delta25}$ heterozygous adults and offspring raised to adulthood. Heterozygous F1 *lamb1a*; *lamb1b* adults were genotyped using the relevant primers for each locus and used for experiments.

*Quantification of heart rate*

Prior to imaging at 2dpf, embryos were sorted based on morphology into siblings and mutants. A pair of embryos (1 sibling, 1 mutant) were transferred in E3 medium from a 28.5oC incubator. A single embryo was positioned laterally on an agarose mould (2% agarose in E3) for imaging under a dissection microscope (11.5X magnification) attached to a High Speed Camera (Chameleon3 USB3, FLIR Integrated Imaging Solutions Inc) focussed on the heart. Image sequences (.tif) of 5s were captured at 150 frames per second using SpinView Software (Spinnaker v. 2.0.0.147). This procedure was repeated for the remaining embryo of the pair and was then repeated at 3dpf. Image sequences were imported into Fiji and converted to .avi movies. Movies were imported into and heart rate quantified in DanioScope (Noldus). Individual values represent an average heart rate over the 5s imaging period.
